# SCITRAM: a Single-cell Integrated transcription regulation modeler for predicting master transcription factors

**DOI:** 10.64898/2026.09.30.755779

**Authors:** Ariel Galindo-Albarran, Maximilien Duvina, Paula Barbao, Alba Rodriguez-Garcia, Sonia Guedan, Christoph Heuser-Loy, Luca Gattinoni, Marco Antonio Mendoza-Parra

## Abstract

Single-cell omics technologies have provided unprecedented access to molecular cell heterogeneity, revolutionizing our understanding of complex living systems. Although generating such type of data has become a streamlined process, data processing still relies on command-line driven tools, or commercial solutions. Furthermore, single-cell transcriptomics analysis is usually reduced to stratifying cells on the grounds of gene expression signatures, while deconvolving gene regulatory programs (and their associated master transcription factors) responsible for cell heterogeneity are not classically addressed with the available tools, or performed under different data processing platforms.

Herein, we present SCITRAM (Single-Cell Integrated TRAnscription regulation Modeler), a user-friendly stand-alone computational solution for predicting master transcription factors from single-cell transcriptomics data. SCITRAM first reconstructs a primary gene regulatory network and subsequently uses this network as template to model transcriptional cascades driven by the in-silico activation of transcription factors identified within the system.

We validated SCITRAM performance across a variety of cell fate transition events including T cell exhaustion during solid tumor treatment, lineage stratification during pancreatic endocrinogenesis, as well as nervous tissue formation in cerebral organoid models. Overall, SCITRAM provides an accessible and integrated platform for processing (large amounts of) single-cell transcriptomics data, enabling the investigation of transcriptional regulatory programs and their master transcription factors without requiring a command-line driven environment.

**Key Points:**

- **SCITRAM fills a methodological gap** by providing an integrated, user-friendly platform for inferring gene regulatory programs and master transcription factors from single-cell transcriptomics data, beyond classical clustering-based analyses.
- **Its workflow combines GRN reconstruction with in-silico TF activation modeling,** enabling systematic exploration of transcriptional cascades underlying cellular heterogeneity.
- The tool eliminates reliance on command-line pipelines or commercial software, making large-scale single-cell regulatory analysis accessible to non-computational users.
- **SCITRAM was validated across diverse biological systems,** including T-cell exhaustion, pancreatic endocrine lineage specification, and neural tissue formation in cerebral organoids.
- **SCITRAM provides a unified environment** for processing and interpreting single-cell transcriptomics data with a focus on regulatory program discovery.

**Biographical Note:** Marco Antonio Mendoza-Parra; PhD/HDR. Research Director (CNRS – France); Head of the team Systems Biology of cell fate decisions (SysFate: https://www.sysfate.org/).

## INTRODUCTION

Complex living systems are structured by an assembly of a variety of cells with diverse molecular and functional characteristics, which, in addition, can change their fate over time and in response to their spatial and cellular microenvironment. Importantly, cell fate and their inherent transitions into novel states are driven by defined gene regulatory programs controlled by groups of transcription factors acting together in a coordinated manner. Accessing to gene expression changes defining cell heterogeneity has been strongly accelerated with the arrival of single-cell omics technologies. Indeed, nowadays the assessment of transcriptome profiling over several thousand of single-cells has become a streamlined process for interrogating cell heterogeneity in blood and, following dissociation, for dissecting cellular complexity of organs and tissues.

While accessing to gene expression at a single-cell level can be performed under rather well stablished conditions, the required data processing for extracting meaningful information relies in a variety of computational tools, mostly available in command-line driven environments^1^. Furthermore, single-cell transcriptomics data analysis is mainly focused on stratification and cell-type annotation based on gene expression signatures, while aspects concerning the unveiling of the various gene regulatory programs defining cell heterogeneity are not typically addressed, or performed under different data processing platforms and multiple command-line scripts that need to be run sequentially ^2–4^.

Herein we introduce SCITRAM, a user-friendly computational solution able to (i) process multiple single-cell transcriptomics files at once, (ii) perform dimensionality reduction while retaining the origin of cells retrieved on the obtained clusters, (iii) compute differential gene expression profiling among stratified clusters; (iv) annotate clusters based on user-provided cell-type gene association information; (v) reconstitute gene regulatory networks per cluster by using transcription factors-target gene relationships; (vi) classify transcription factors as master drivers defining cluster differences retrieved within single-cell transcriptomics readouts; and (vii) display transcription factor-coregulatory networks to reveal their level hierarchy.

This strategy, which has been initially described in our previous computational tool TETRAMER for the analysis of bulk transcriptomics data^5,6^, has been validated here across a variety of datasets, including the analysis of CAR-T cells during solid tumors treatment, lineage stratification detection during pancreatic endocrinogenesis, as well as nervous tissue formation in cerebral organoid models.

## RESULTS

SCITRAM has been designed for processing single-cell transcriptomic readouts generated by most common strategies. Notably, primary processed files in either h5 or in matrix/barcode/feature three files format are required as input. Single-cell transcriptomics data uploaded in SCITRAM is first stratified by applying non-linear dimensionality reduction technique UMAP (Uniform Manifold Approximation and Projection) (**Figure 1**). Stratified cells are used for assessing differential gene expression levels between selected clusters. Differential gene expression levels can be visualized by MvA plots (minus vs average), as well as by Volcano plots, for which an adjusted P-value is computed by comparing both log2 fold-change (LFC) distributions associated to the compared clusters. The obtained differential gene expression readouts are integrated with a collection of prior knowledge Transcription factor (TF) – target gene (TG) regulation relationships^7^, leading to the construction of a gene regulatory network (GRN). Considering that this integrative effort could introduce false TF-TG relationships, a machine driven strategy based on the in-silico activation of all TFs (one by one) and the tracing of the transcription regulation cascades issued from such activation is used for clearing (i) TF-TG relationships (edges) that do not lead to the activation/repression of a downstream target; (ii) edges incoherent with the discretized differential gene expression status that are part of the connectivity;(iii) edge connections that are against the downstream flux of the information (**Figure 1**). This machine-driven strategy allows to reduce the GRN complexity towards a connectivity in coherence with the differential gene expression information issued from the single-cell transcriptomics analysis. With the aim of identifying the most influent TFs within the system, TFs are ranked based on the fraction of their downstream targets (direct and indirect) relative to the total number of genes within the reconstructed GRN. This metrics, defined as the TF master regulatory index (MRI), allows to highlight the most influential TFs. Furthermore, a randomization of the edges composing the GRN is used for identifying the confidence associated to the inferred MRIs. Finally, a gene co-regulatory network among all significant master TFs is generated, leading to the identification of hierarchy relationships of interest for instance when perturbation assays are aimed to do by experimentalists (**Figure 1**).

**Figure 1.**
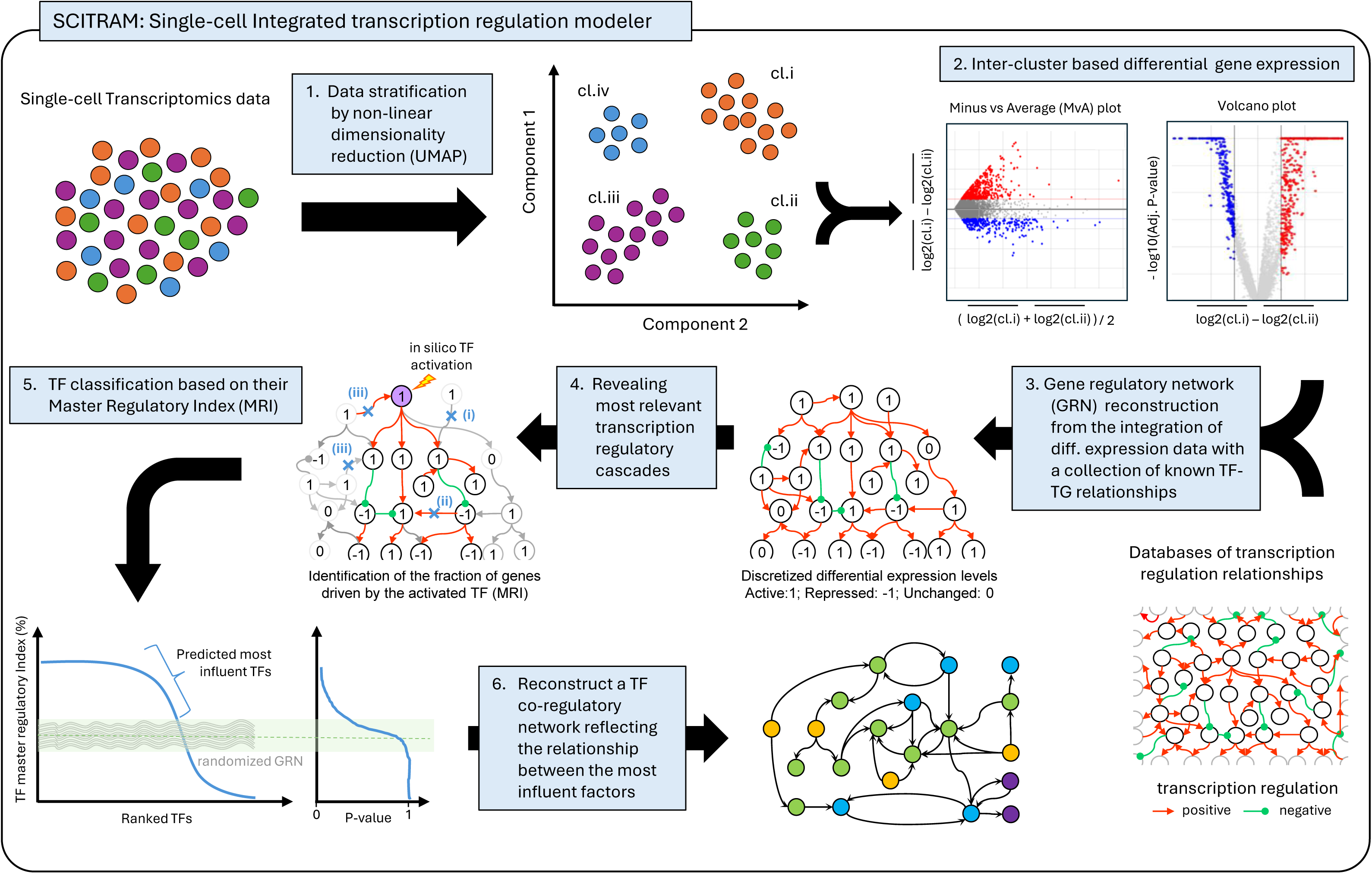
Scheme illustrating the main steps behind SCITRAM. **(1)** Single-cell transcriptomics data uploaded in SCITRAM is first stratified by applying non-linear dimensionality reduction technique UMAP (Uniform Manifold Approximation and Projection). **(2)** Stratified cells are used for assessing differential gene expression levels between selected clusters. Differential gene expression levels can be visualized as MvA plots (minus vs average), as well as Volcano plots, for which an adjusted P-value is computed by comparing both log2 fold-change distributions associated to the compared clusters. **(3)** The obtained differential gene expression readouts are integrated with a collection of prior knowledge Transcription factor (TF) – target gene (TG) regulation relationships, leading to the construction of a gene regulatory network (GRN). **(4)** Considering that this integrative effort could introduce false TF-TG relationships, a machine-driven strategy based on the *in-silico* activation of all TFs (one by one) and the tracing of the transcription regulation cascades issued from such activation is used for clearing (i) TF-TG relationships (edges) that do not lead to the activation/repression of a downstream targets; (ii) edges incoherent with the discretized differential gene expression status that are part of the connectivity;(iii) edge connections that are against the downstream flux of the information. **(5)** The most influent TFs within the system are traced by their ranking based on the fraction of their downstream targets (direct and indirect), relative to the total number of genes within the reconstructed GRN (master regulatory index (MRI)). Furthermore, a randomization of the edges composing the GRN is used for identifying the confidence associated to the inferred MRIs. **(6)** Finally, a gene co-regulatory network among all significant master TFs is generated, leading to the identification of hierarchy relationships.

SCITRAM identifies most relevant universal transcription factors deregulated during T cell exhaustion assessed on pan-cancer single-cell atlas

Immunotherapies are revolutionizing cancer treatment, but discrepancies in performance between cancer types remain a challenge to overcome. Indeed, while tumor-infiltrating T cells are able to kill cancer cells during the early phases of the immune response, over time they enter into a dysfunctional state, known as “T cell exhaustion”. Importantly, such exhaustion state has been shown to be accompanied by the deregulation of a large panel of genes, including multiple TFs. By composing a pan-cancer single-cell atlas covering > 397 thousands of T cells issued from 21 cancer types obtained from 316 patients, Zheng and colleagues established a universal collection of master TFs driving transcriptional regulation of CD8+ T cell exhaustion ^8^ (**Figure 2A**). To evaluate the performance of SCITRAM in capturing master TFs involved in T cell exhaustion, we have reanalysed the single-cell data collection provided by Zheng and colleagues. Dimensionality reduction analysis (UMAP) performed by SCITRAM over > 397 thousand cells revealed 17 clusters (**Figure 2A**). By using a panel of genes associated to either the effector or the exhausted state, we have identified specific cell populations (**Figure 2B**), that could be clearly associated with distinct clusters within the UMAP. Indeed, as illustrated in **Figure 2C** and **Supplementary Figure S1**, clusters 2 & 15 within the UMAP are enriched in genes associated to effector cells, while cluster 14, 6, 8 and 10 are preferentially enriched in markers associated to exhaustion.

**Figure 2.**
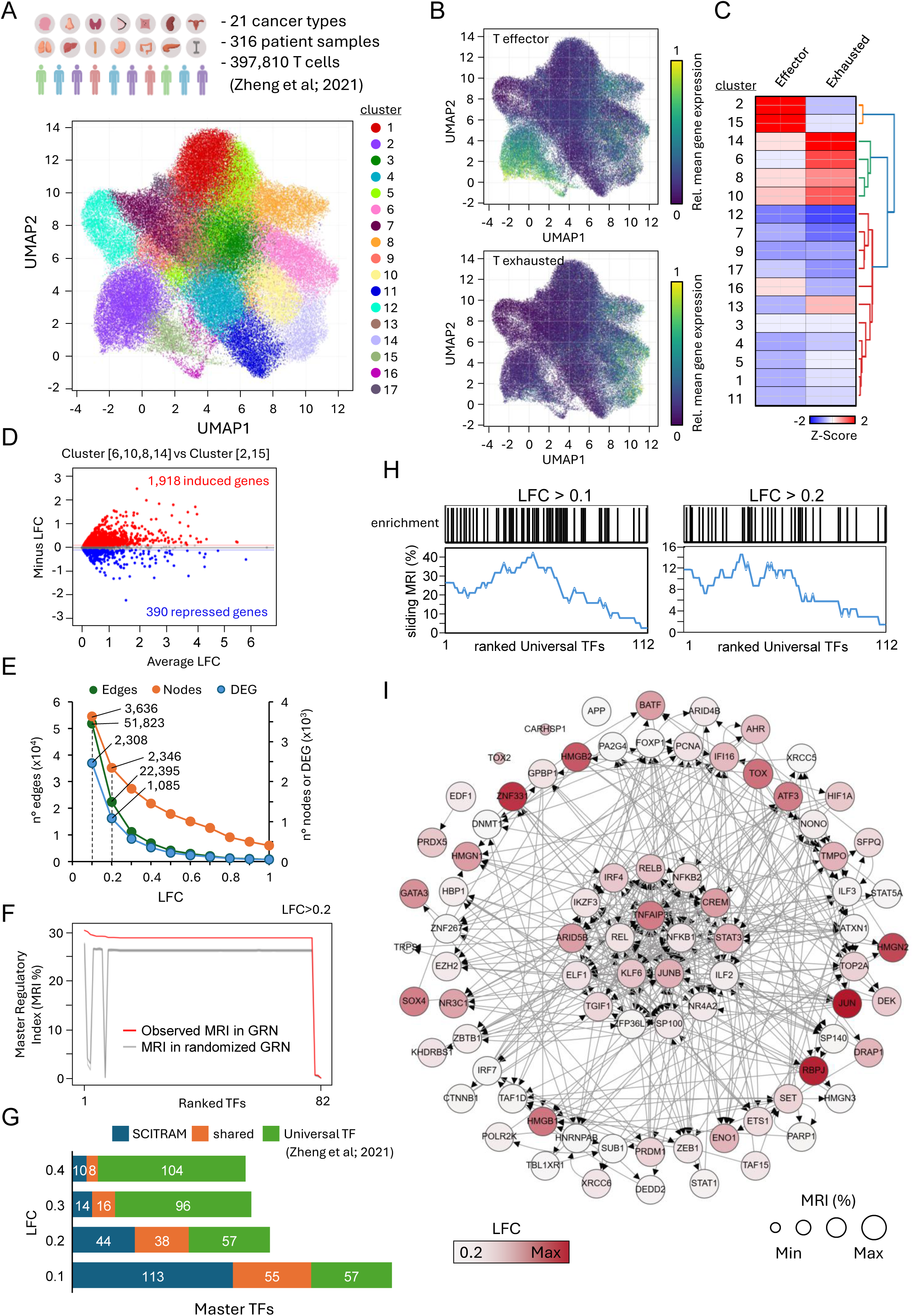
SCITRAM captures most relevant universal TFs deregulated during T cell exhaustion assessed on pan-cancer single-cell atlas and reveal novel players. (A) Top: Scheme describing the complexity of the pan-cancer single-cell atlas constituted by Zheng et al. Bottom: Dimensionality reduction analysis (UMAP) performed by SCITRAM over >397 thousand T cells composing the pan-cancer single-cell atlas. (B) Cells presenting high enrichment levels of gene markers associated to the effector (top panel), or exhausted state (bottom panel). Heatmap colour scale corresponds to the relative mean gene expression. (C) Normalized z-score gene expression per cluster associated to the effector or the exhausted state. (D) Minus vs Average (MvA) gene expression plot displaying the comparison between clusters 6,10,8,14 (exhausted) and cluster 2&15 (effector). Induced (red) and repressed (blue) genes are defined by a log2 fold-change (LFC) filter of 0.1. (E) Number of nodes (genes; right Y-axis, orange) and edges (TF-TG relationships; left Y-axis, green) retrieved in the gene regulatory network reconstructed by SCITRAM by using differentially expressed genes (DEG; right Y-axis, blue) obtained at different LFC thresholds (X-axis). Dashed lines highlight GRNs built for a LFC threshold of 0.1 and 0.2. (F) Transcription factors ranked based on their master regulatory index inferred from the reconstructed GRN (LFC>0.2). (G) Comparison between the list of TFs associated with exhaustion defined by Zheng et al. (Universal TF) and those obtained by SCITRAM thanks to the computed master regulatory index. Shared TFs between both strategies are displayed in orange, while SCITRAM or Universal TF specific TFs are displayed in blue or green respectively. (H) Top: Enrichment of the 112 Universal TFs presented by Zheng et al and displayed based on their raking metrics. Black bars indicate the presence of the universal TF within the SCITRAM analysis; while white regions reflect the absence of the such TFs. Bottom: Mean MRI computed over 20 ranked TF readouts and slided (1 position) over the whole universal TF list. Displays were performed for both LFC>0.1 and LFC>0.2. (I) Gene co-regulatory network displaying the relationship between all master TFs retrieved by SCITRAM (LFC>0.2).

To reveal deregulated genes between exhausted and effector cells, we used SCITRAM to perform a differential expression analysis between the aforementioned clusters. This analysis led to the identification of more than 1,900 genes that were induced in exhausted cells, while an additional 390 genes were significantly downregulated (LFC threshold: 0.1; **Figure 2D**). This 2,308 differentially expressed genes (DEGs) were subsequently used to build a gene regulatory network, composed of 3,636 nodes and more than 51 thousand TF-TG relationships (edges) (**Figure 2E**). SCITRAM also provides the possibility to reconstruct GRNs at various LFCs in an automated manner (by defining LFC intervals and evaluation steps) enabling to assess the influence of the LFC thresholding parameter for the master TFs identification. As illustrated in **Figure 2E**, a direct correlation between the number of DEGs and the size of the GRN (defined by the number of nodes and edges) can be obtained from such automated processing, allowing to select the best LFC conditions for the downstream analyses. For instance, for an LFC>0.2; a smaller GRN has been obtained, composed by 2,346 nodes and ∼22 thousand edges (**Figure 2E**). Furthermore, with this LFC threshold conditions, we found a total of 82 TFs with a significant master regulatory index (**Figure 2F**). Notably, 38 of these TFs are part of the universal TFs proposed by Zheng and colleagues (**Figure 2G & Supplementary Figure S2**). By relaxing the LFC threshold conditions to 0.1 a total of 55 “Universal TFs” were detected (**Figure 2G & Supplementary Figure S2**); which in addition correspond to the top-ranked factors based on the criteria defined by Zheng and colleagues (**Figure 2H**) (significantly high expression in >80% of analyzed cancers ^8^).

A gene co-regulatory network generated from the master TFs predicted by SCITRAM reveals the presence of factors like TOX, TOX2, RBPJ, PRDM1, VDR, IKZF4, STAT3, IFI16; described as the top-10 within the Universal TF list; as well as SOX4, ETV1, BHLHE40 or IKZF3; also retrieved within this study as major TFs ^8^. In addition, among the 44 TFs found specifically by SCITRAM we can highlight EDF1, which is known to act as a critical metabolic and transcriptional coactivator in tumor cells,driving CD8+ T cell dysfunction and exhaustion^9^. Furthermore, the Activating Transcription Factor 3 (ATF3), known to drive immunosuppression and promote T cell dysfunction when it is over-expressed in the tumor microenvironment ^10^, has been also predicted as master TF.

Other factors like the High Mobility Group Box 2 (HMGB2); known to be induced in exhausted CD8+ T cells during chronic viral infections and cancer ^11^; or TNFAIP3; known to act as a critical negative regulator of NF-κB signaling and a key driver of T cell dysfunction and exhaustion in cancer environments like acute myeloid leukemia (AML) and solid tumors ^12^; were also found as a key master TF specifically predicted by SCITRAM.

In summary, SCITRAM allowed to process >397 thousand T cell readouts, assessed from the analysis performed by Zheng and colleagues, in a rather straight manner, leading to the identification of master TFs of the T cell exhaustion process. This analysis covered cells stratification, clusters annotation thanks to the upload of a cell types annotation file (62 gene markers interrogated at once; **Supplementary Table S1**), differential expression analysis between selected clusters, GRNs construction for different LFC thresholds, master TFs prediction, and master TF co-regulatory maps display.

SCITRAM captures master transcription factors driving exhaustion during solid tumor treatment with CAR-T cells

Chimeric antigen receptor (CAR)-T cell therapy has revolutionized the treatment of hematologic tumors, reaching remarkable clinical results including complete remission in patients with certain blood cancers. Due to this major clinical success, several CAR-T treatments have been approved by the FDA (US Food and Drug Administration). Unfortunately, such achievements remain still to be improved in the case of solid tumors, notably also due to the immunosuppressive tumor microenvironment leading to CAR-T cell exhaustion ^13^.

To identify key master TFs that are deregulated during CAR-T cell exhaustion in solid tumor treatment, we have analyzed single CD8+ T cell transcriptomics data generated from an ovarian xenograft tumor model (SKOV3) treated with HER-2 CAR-T cells ^14^. To capture the transcriptional dynamics associated with CAR-T cell activation and subsequent exhaustion, tumor-infiltrating T cells were isolated at three distinct stages of therapy: prior to CAR-T cell infusion (pre-infusion; day 0), following CAR-T cell infusion during the effector phase (day 8), and after therapy failure during the exhausted phase (day 27) (**Figure 3A**). These stages were defined based on tumor growth kinetics monitored throughout CAR-T cell treatment (**Figure 3B**).

**Figure 3.**
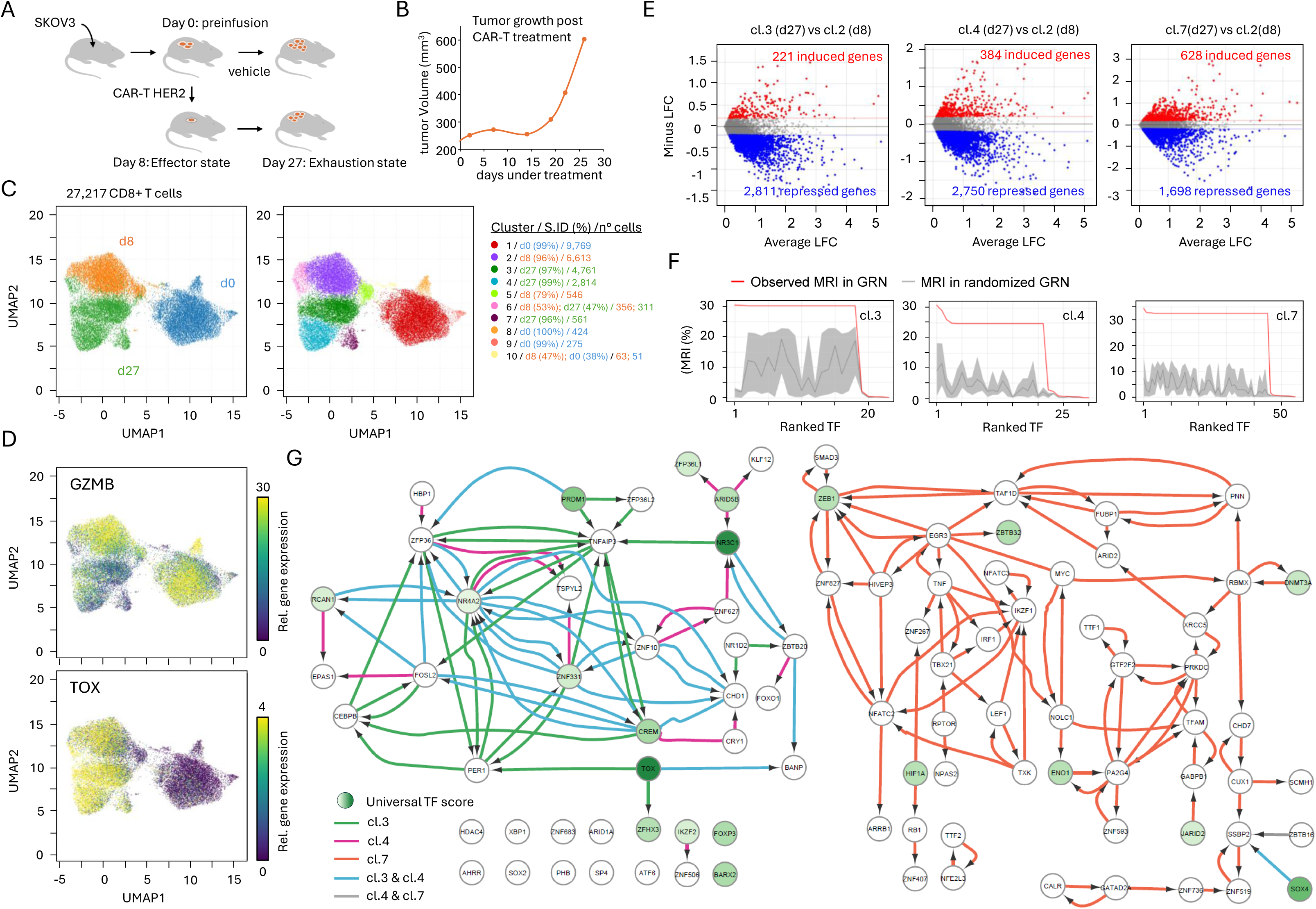
SCITRAM captures master transcription factors driving exhaustion during solid tumor treatment with CAR-T cells. **(A)** Scheme illustrating the ovarian xenograft tumor model (SKOV3) in which CAR-T cells targeting the HER2 protein are used for killing the tumor. **(B)** Tumor growth measurements assessed post CAR-T treatment. **(C)** Dimensionality reduction analysis (UMAP) performed by SCITRAM over >27,000 CD8+ T cells issued from three temporal stages; day 0 (preinfusion); day 8 (effector); and day 27 (exhaustion) (Left panel). Cells were clustered based on their gene expression similarities in 10 different clusters; for which the origin of the most enriched fraction (S.ID: sample Identifier) and number of cells per cluster is indicated (right panel). **(D)** Gene expression associated either to the effector marker Granzyme-Beta (GZMB) or the exhaustion marker TOX depicted over the UMAP dimensionality reduction map. **(E)** Minus versus Average plot displaying the differential gene expression analysis between clusters associated to the exhaustion state (day 27: cl.3; cl.4 & cl.7) and cluster 2 enriched in cells issued from the effector state (day 8). Gray zone corresponds to non-differentially expressed genes. Log2 fold-change (LFC) threshold: 0.2. **(F)** Master regulatory Index associated to the ranked TFs computed by SCITRAM over gene regulatory networks reconstructed from the differentially expressed genes displayed in panel (E). **(G)** Gene co-regulatory network associated to the master TFs obtained in (F). Edge colours indicate the cluster in which the master TF has been found. Node gradient colour correspond to the “Universal TF scoring” proposed by Zheng et al^8^.

Single-cell Transcriptomics assay performed on 27,217 CD8+ T cells was processed with SCITRAM, revealing a clear separation between cells issued from the pre-infusion state (day 0); and those isolated from either the effector (day 8) or exhaustion phase (day 27). These three major temporal states were further stratified in 10 cell clusters based on their gene expression similarities (**Figure 3C**). The loss on performance of T cells over time was confirmed by the decrease in expression levels observed for the effector marker GZMB (Granzyme B); as well as their transition into an exhausted state, revealed by the over-expression of the exhaustion master TF, TOX (**Figure 3D**).

To further identify gene expression changes occurring during the effector-exhaustion T cell transition, we performed a differential gene expression analysis between cell clusters associated to each of these two states. We found that cluster 3, 4 and 7 were predominantly composed of cells isolated at day 27 (> 95%); while cluster 2 was enriched in cells isolated at day 8 (96%) (**Figure 3C**). A differential expression analysis performed by SCITRAM between the aforementioned clusters revealed 221 up-regulated genes in cluster 3 relative to cluster 2; other 384 up-regulated genes in cluster 4 relative to cluster 2 and 628 up-regulated genes in cluster 7 (**Figure 3E**). Importantly, >60% of the up-regulated genes in cluster 3 were also found in cluster 4, in agreement with their spatial proximity within the UMAP; while cluster 7 presented less than 10% of the up-regulated genes shared with either cluster 3 or 4 (**Supplementary Figure S3**).

By using the differentially expressed genes retrieved between cluster 3,4 & 7 relative to cluster 2, we reconstructed GRNs composed by 3,884 nodes and 26,150 thousand TF-TG relationships (edges) for cluster 3; 3,925 nodes and 28,135 edges for cluster 4; 4,359 nodes and 30,593 edges for cluster 7 (**Supplementary Figure S4**). These three GRNs were then used for retrieving master TFs thanks to their master regulatory index inferred by SCITRAM (**Figure 3F**). By integrating master TFs information retrieved over all three clusters, we identified 26 TF-TG relationships specific to cluster 3; 16 to cluster 4; but also 28 others retrieved in both clusters (**Figure 3G**). Similarly, 94 edges were found specifically associated to cluster 7, consistent with its greater transcriptional distance from cluster 2 compared with clusters 3 and 4. Twenty of the TFs identified in our analysis were also included in the Universal TF list described by Zheng and colleagues. These included established regulators of T-cell function and differentiation, such as TOX, NR3C1, FOXP3, BARX2, IKZF2, ZFHX3, CREM, PRDM1, NR4A2, RCAN1, ARID5B, ZNF331 or ZFP36L1; all of them retrieved either in cluster 3 or cluster 4. Conversely, Factors such as SOX4, JARID2, ENO1, HIF1A, DNMT3A, ZBTB32 or ZEB1 were specifically enriched in cluster 7. Notably, master TFs identification performed by SCITRAM clearly identified early exhausted (cluster 3,4) and terminally exhausted players, in agreement with previous classification^15–17^.

Lineage stratification detection during pancreatic endocrinogenesis performed by master gene co-regulatory networks reconstruction.

Single-cell transcriptomics is a powerful strategy for resolving cellular heterogeneity and defining lineage specialization at high resolution. In particular, tracing the gene regulatory programs that govern transitions from a progenitor status to more specialized cell systems requires to reconstruct the underlying gene regulatory networks from the information embedded within single-cell transcriptomics data.

Using a pancreatic endocrine development single-cell transcriptomics data^18^, Wang W. and colleagues applied RegVelo, a computational tool able to infer gene regulatory networks from single-cell transcriptomes, to recover cell lineage formation^2^. To assess the performance of SCITRAM in comparison with RegVelo in this cell fate specialization setting, we analyzed the same pancreatic endocrinogenesis dataset, identifying 12 distinct clusters covering a total of 3,658 cells (**Figure 4A**). Using pancreatic cell type gene markers available in the Panglao Database ^19^ (**Supplementary Table S1**), we have annotated the identified pancreatic clusters as corresponding to (i) progenitor cells, (ii) epithelial lining ductal cells, as well as (iii) hormone-producing endocrine Alpha, Beta, Delta or Epsilon cells (**Figure 4B and Supplementary Figure S5**). Furthermore, the enrichment display of these different cell type markers within the stratified cells revealed a clear separation between ductal and hormone-producing endocrine cells (**Figure 4C**), also visible in the mean marker gene expression enrichment levels per cluster (Z-score normalized), as displayed in **Figure 4B**.

**Figure 4.**
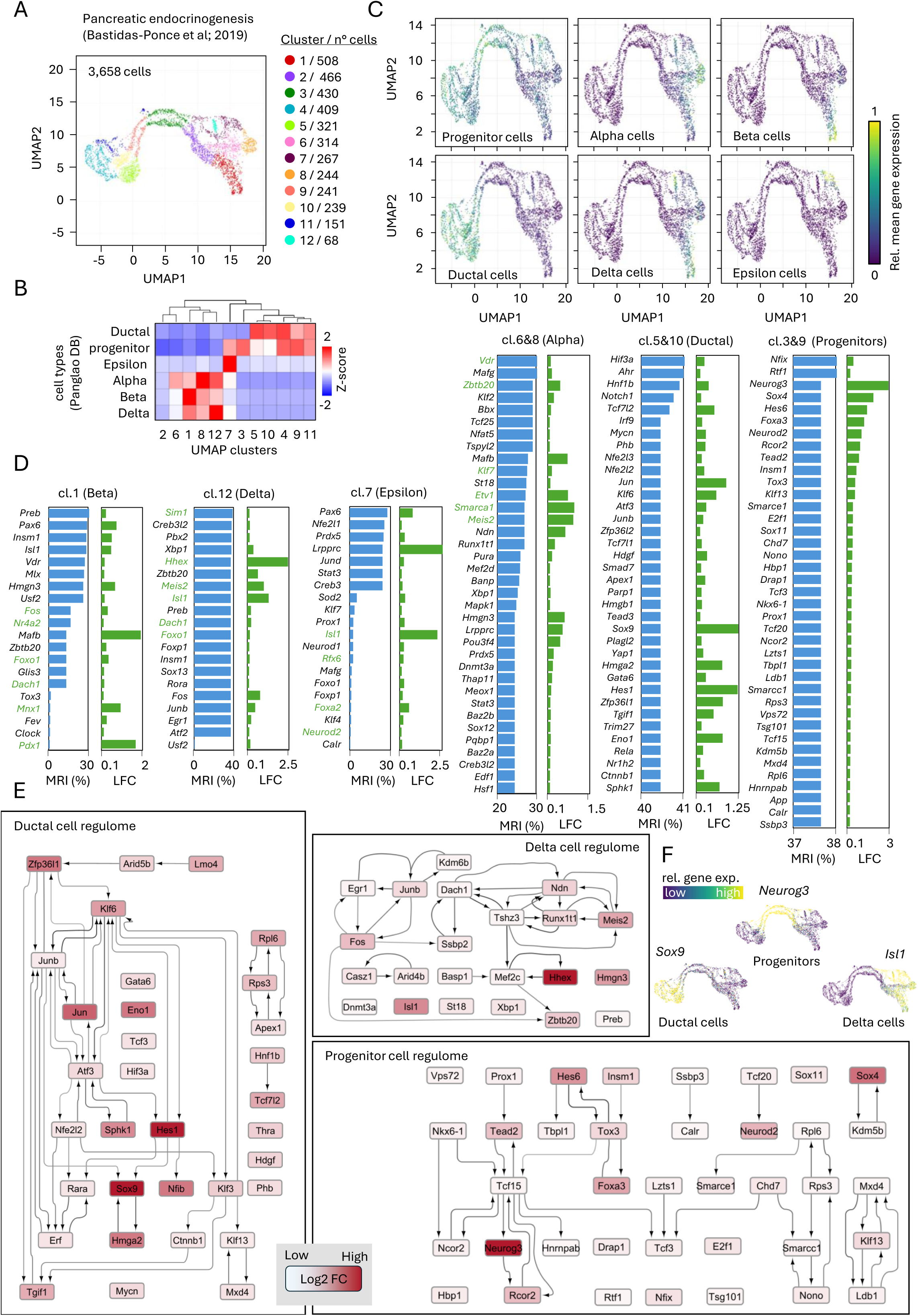
Master transcription drivers involved in pancreatic endocrinogenesis as revealed by SCITRAM. **(A)** Dimensionality reduction analysis (UMAP) performed by SCITRAM on pancreatic endocrinogenesis data generated by Bastidas-Ponce et al^18^. **(B)** Heatmap displaying the enrichment of gene markers (reported by Pangalo Database) associated to the indicated pancreatic cell types per cell clusters revealed in (A). **(C)** Relative mean gene expression associated to the markers per indicated cell types displayed over the UMAP stratification view. **(D)** Ranked master TFs per cluster associated to the indicated pancreatic cell types (blue) and their corresponding fold-change (log2) (green) relative to all other clusters. MRI: master regulatory index. LFC: Log2 fold-change. Gene names in green correspond to TFs found as part of the top 20 putative drivers as predicted by RegVelo^2^. **(E)** Master TF co-regulatory networks assessed for the indicated pancreatic cell types. Displayed GRNs were been performed for a log2 fold-change threshold of 0.2, as a way to reduce network complexity. **(F)** Relative gene expression levels displayed on the UMAP view of the indicated master TFs associated to the cell type regulomes.

To infer master TFs driving the fate of each of the indicated cell types, we have reconstructed GRNs. For this, we first compared clusters 6&8 with all other clusters to build a GRN associated to the hormone-producing endocrine Alpha cells. This analysis revealed a total of 3,494 differentially expressed genes, which gave rise to a GRN composed of 4,379 nodes and more than 40 thousand edges (**Supplementary Figure S6**). By modeling transcription regulation propagation over the reconstructed GRN we identified 36 TFs with a master regulatory index >24% (**Figure 4D**). Among them, several of the putative drivers predicted by RegVelo such as *Vdr*, *Zbtb20*, *Klf7*, *Etv1*, *Smarca1*, *Meis2* were also captured by SCITRAM. Furthermore, other factors like *Mafb*, a crucial TF that controls the development, maintenance and function of pancreatic alpha cells, notably by being critical for glucagon production and secretion^20^, was uniquely predicted by SCITRAM as a master TF (**Figure 4D**). A similar analysis performed for the remaining hormone-producing endocrine cells inferred a total of 15 TFs (MRI> 13%) for the Beta cells; 19 TFs (MRI>37%) for the Delta cells; and 7 TFs (MRI>24%) for the Epsilon cells, respectively (**Figure 4D**). Importantly, while several of the putative drivers predicted by RegVelo, including *Sim1*, *Hhex*, *Meis2*, *Isl1*, *Dach1*, or *Foxo1* presented relatively high MRI levels for Delta cells, only few of them like *Fos*, *Nr4a2*, *Foxo1* or *Dach1* appeared as master regulators for Beta cells (**Figure 4D**). Furthermore, none of the factors predicted by RegVelo as their top-20 putative drivers for the Epsilon cells presented significant MRIs in our analysis (**Figure 4D**). In contrary, SCITRAM predicted *Pax6*, *Insm1*, or *Hmgn3* as master TFs in Beta cells; in agreement with previous studies describing their major role in the maintenance of Beta cell identity^21–23^.

In a similar manner, we have compared clusters 3&9 with all other clusters to build a GRN associated to the pancreatic progenitor cells, or clusters 5&10 to reconstruct a GRN linked to ductal cells. By modeling transcription regulation propagation over the reconstructed GRNs we identified 39 TFs (MRI >37%) in pancreatic progenitors, and 36 TFs (MRI>40%) in ductal cells. By reconstructing a TF co-regulatory network, SCITRAM allowed to highlight the main role of several TFs, known to participate at the pancreatic progenitor state, including *Neurog3*, *Tcf15*, *Neurod2*, or *Sox4* (**Figure 4E**). Similarly, the master TF-coregulatory network reconstituted for Ductal cells reveal the connectivity between major players like *Hes1*, *Sox9*, *Klf6*, *Tgif1* or *Hnf1b* (**Figure 4E**). Finally, the reconstituted master TF regulome for the hormone-producing endocrine Delta cells revealed the transcription regulation between factors like *Hhex* and *Mef2c*; both known to control the molecular identity and development of pancreatic Delta cells; the hub nature of *Runx1t1*, known to be involved in the lineage trajectory of progenitor cells that give rise to hormone-producing islet cells, including somatostatin-secreting delta cells^24^; or the presence of *Isl1*, a major TF for the survival and identity of pancreatic Delta cells^25^ (**Figure 4E**). In all cases, master TF expression within the stratified single-cell transcriptomics data also confirmed their regional presence, further supporting the capacity of SCITRAM to properly capture master TF players during cell lineage specialization processes like pancreatic endocrinogenesis (**Figure 4F & Supplementary S6-S8**).

Fate specification programs of the human cerebral cortex traced by SCITRAM during brain organoids progression.

While the study of nervous tissue formation and brain development traditionally relied on animal models, the advent of three-dimensional, self-organized cerebral organoid cultures derived from human induced pluripotent stem cells, has provided an *in vitro* model for investigating the cellular and molecular mechanisms underlying these cell fate decisions in a human setting. By combining cerebral organoid cultures with single-cell transcriptomics profiling, Uzquiano and colleagues traced nervous tissue formation collecting samples at different time-points covering a period of 6 months^26^ (**Figure 5A**). To evaluate the performance of SCITRAM in identifying master transcriptional events driving cell fate decisions during brain organoids progression, we processed the single-cell transcriptomics data generated by Uzquiano et al encompassing 8 different time-points for a total of 302,242 cells. SCITRAM stratified cells in 18 clusters, which, in addition, presented a UMAP spatial organization coherent with the temporal transition during brain organoid progression (**Figure 5B**). Indeed, a hierarchical clustering applied to the number of cells per cluster and time-point of origine clearly subdivided the events in two major groups, namely clusters associated to early stages (23 days, 1,1.5 and 2 months of culture) and those involved in the late organoids’ progression (3, 4, 5 and 6 months) (**Figure 5C**). Based on this classification, we used SCITRAM for revealing master TFs associated to clusters retrieved either in the early or late phase of brain organoids’ progression. We found 48 master TFs associated with cluster 3, primary composed of cells issued from the early stages of organoids differentiation. Among them, we can highlight the LIM homeobox 2 transcription factor (LHX2), known to play a critical role during neural development by promoting the expression of PAX6 and by repressing non-neural differentiation pathways such as BMP and WNT signaling^27^. Cluster 3 was also enriched in PAX6, SOX2, and DACH1, which are critical for maintaining neural progenitor identity^28,29^, and HES5, known to inhibit neuronal differentiation and maintain neural stem and progenitor cells in a dividing state ^30^ (**Figure 5D**). A gene ontology (GO) terms enrichment analysis performed on the 48 master TFs enriched in cluster 3 revealed the presence of signatures related to radial glial cells, neural progenitors and neural stem cells, in agreement with an early state during nervous tissue differentiation (**Figure 5E**).

**Figure 5.**
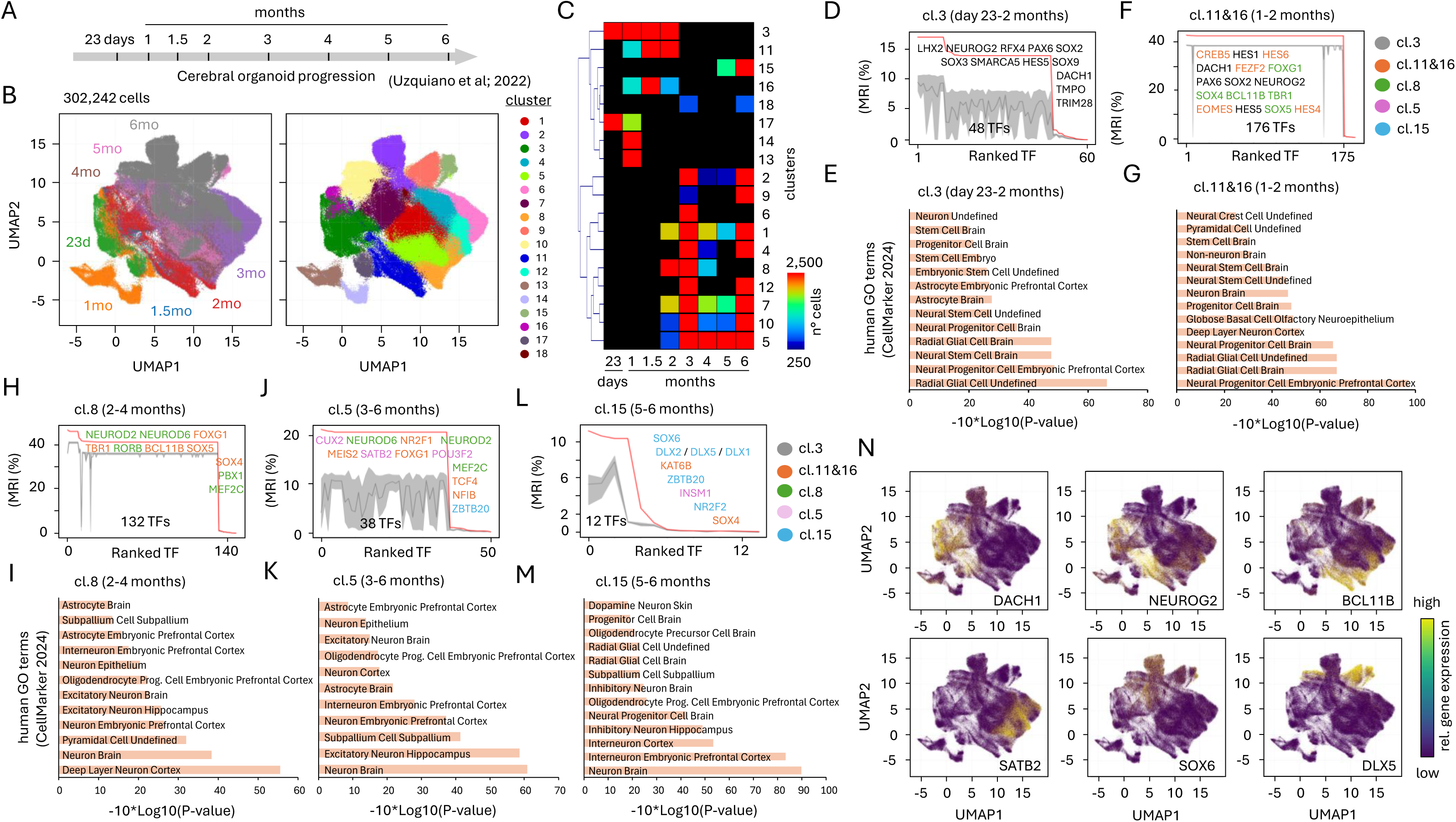
Fate specification programs of the human cerebral cortex traced by SCITRAM during brain organoids progression. **(A)** Scheme illustrating the time-frame used by Uzquiano et al. to collect samples for single-cell transcriptomics data during human cerebral organoids progression. **(B)** Dimensionality reduction analysis (UMAP) performed by SCITRAM over >300 thousand dissociated cells issued from all indicated time-points in (A). Left panel: UMAP stratification colored based on the collected time-points. Mo= months; 23d= 23 days. Right panel: Cells clustered on 18 groups based on their gene expression similarities. **(C)** Heatmap displaying the number of cells per cluster issued from the different organoid time-point samples (Hierarchical clustered). **(D)** Ranked TFs based on their master regulatory index (MRI) associated to the differential expression analysis of cluster 3 (enriched in cells issued from day 23-2months) relative to all other clusters. Red line: MRI corresponding to the top 60 TFs. Gray zone: MRI generated by SCITRAM by randomizing all edges composing the reconstructed GRN. Gray zone is produced by 20 randomization processes. Gray line corresponds to the mean MRI obtained from the 20 randomized processes. Example of top-ranked master TFs are listed. **(E)** Human Gene Ontology (GO) enrichment analysis (CellMarker Database) performed on the 48 master TFs retrieved in (D). **(F)** Same as (D) but performed for clusters 11&16 (enriched in cells issued from 1-2months) relative to all other clusters. Colored master TFs correspond to clusters in which they are also observed. **(G)** Same as (E) but performed for the 176 master TFs found in clusters 11&16. **(H)** Same as (F) but performed for cluster 8 (enriched in cells issued from 2-4months) relative to all other clusters. **(I)** Same as (G) but performed for the 132 master TFs found in cluster 8. **(J)** Same as (H) but performed for cluster 5 (enriched in cells issued from 3-6months) relative to all other clusters. **(K)** Same as (I) but performed for the 38 master TFs found in cluster 5. **(L)** Same as (J) but performed for cluster 15 (enriched in cells issued from 5-6months) relative to all other clusters. **(M)** Same as (K) but performed for the 12 master TFs found in cluster 15. **(N)** Relative gene expression display within the UMAP stratification map associated to the indicated master TF genes, over-expressed at different stages during human cerebral organoids differentiation.

To further identifying TFs driving the progress in nervous tissue development we interrogated the cell fate associated to clusters 11&16, composed by cells collected between 1- and 2-months during organoids progression. In addition to recovering 42 of the 48 TFs found in cluster 3, SCITRAM identified 134 additional master TFs in these two clusters (**Supplementary Figure S9**), among them CREB5 (known to play a critical role in neuronal lineage specification, stem cell proliferation and axonal survival), HES6 (known to promote neurogenesis and inhibit astrocyte emergence^31^), EOMES/TBR2 (known to be transiently expressed in cortical progenitor cells and to regulate neurogenesis^32^), or HES4 (a transcriptional repressor maintaining neural stem cells in their undifferentiated, proliferative state^33^) (**Figure 5F**). The GO term analysis performed over the 176 master TFs enriched in clusters 11&16 appeared quite similar to those observed in cluster 3, but also included novel cell type signatures related to deep layer neural cortex notably due to the emergence of novel TFs like TBR1, FEZF2, SOX5 or BCL11B/CTIP2 at this stage of differentiation (**Figure 5F&G**).

We next focused on cluster 8 to interrogate cell fate events taking place between 2 and 4 months of organoids progression. SCITRAM revealed a total of 132 master TFs, of which 37 were specifically shared with clusters 11&16 (**Supplementary Figure S9 and Figure 5H**). At this stage, the GO terms enrichment analysis revealed the presence of signatures associated to specialized excitatory neurons, notably due to the presence of master TFs like NEUROD2 and DEUROD6 (known master regulator for postmitotic cortical excitatory neurons^34^), the Retinoic Acid-Related Orphan Receptor Beta (RORB; known as marker of layer 4 cortical neurons in both mouse and human^35^), or MEF2C (known to regulate synapsis activity of specialized neuronal subtypes^36^) (**Figure 5H&I**).

A similar analysis performed on cluster 5, composed by cells issued from 3 to 6 months of brain organoids progression, revealed an enrichment of cell type signatures associated to specialized excitatory neurons, notably associated to the upper cortical layers as confirmed by the presence of the master TFs CUX2 ^37^, SATB2 ^38^, or POU3F2/BRN2 ^39^ (**Figure 5J&K**). Finally, the analysis of cluster 15 (covering cells issued from the 5^th^ and 6^th^ months of organoid differentiation) revealed the presence of cell type signatures associated to cortical interneurons, as supported by the presence of the TF SOX6 (known to control interneurons migration, positioning, diversity and maturation^40^); as well as the emergence of specialized inhibitory neurons, as attested by the expression of the TFs DLX1, DLX2 and DLX5 (known to be essential for the development of GABAergic neurons^41^) (**Figure 5L&M**).

Overall, the analysis performed by SCITRAM allowed to reveal the cascade of master TF players activated in a timely manner to pass from highly dividing neuroepithelial and radial glial cells (e.g. DACH1), into a neuronal differentiation engagement (e.g. NEUROG2), followed by their subtype specification (e.g. BCL11B), neuronal wiring projections (e.g. SATB2), interneurons migration, positioning, diversity and maturation (e.g. SOX6), as well as the emergence of specialized inhibitory neurons (e.g. DLX5) settle into the system for preventing over excitation of the neuronal architecture (**Figure 5N**).

## DISCUSSION

In this study, we present SCITRAM, a novel stand-alone computational solution able to process single-cell transcriptomics data, from the classical cell stratification processing (dimensionality reduction UMAP), passing by the assessment of differential gene expression analysis between user-selected clusters, till the inference of master TFs from the reconstruction of gene regulatory networks followed by a transcription regulation cascade modelling. It is important to mention that, in contrast to other available solutions, SCITRAM provides a user-friendly environment, devoid of requiring to run scripts within a terminal, or using multiple tools available under different platforms. SCITRAM performance has been validated in multiple case studies, including the detection of master TFs involved in T cell exhaustion either within a large pan cancer atlas collection, or within a CAR-T cell mouse model designed to study T cell exhaustion during solid tumor treatment. In these two cases, SCITRAM not only retrieved most of the well described “universal TFs” participating in the T cell exhaustion process, but it also revealed novel players that are worth to explore experimentally in the future.

In a different biological setting, SCITRAM has been challenged to predict master TFs participating at the murine pancreatic endocrinogenesis, by using a dataset recently evaluated by RegVelo, a computational strategy that leverages RNA velocity to provide a dynamic frame for the analysis of gene regulatory networks^2^. While SCITRAM does not incorporate RNA velocity, SCITRAM predicted several of the top-20 ranked putative drivers inferred by RegVelo. Similarly, the analysis of human cerebral organoids progression by single-cell transcriptomics demonstrated that a straight-forward platform like SCITRAM can access to complex temporal readouts in a rather simple manner. Indeed, several of well know key TFs activated during nervous tissue specialization were captured by the transcription regulation cascade strategy used by SCITRAM to classify TFs.

Beyond the demonstrated performance, SCITRAM relies currently on TF-TG prior knowledge collection established on bulk omic assays. A major add-on for the following years would include the usage of single-cell ATAC-seq readouts combined with their corresponding single-cell transcriptomics data for enhancing the pertinence of the annotation of TF-TG relationships. This being said, we believe that accessing to single-cell ATAC-seq would still provide sparse readouts which cannot fully replace prior-knowledge data issued from bulk readouts. For this reason, we consider that a strategy aiming to count with both types of information, followed by our innovative strategy for validating TF-TG relationships pertinence by machine-driven transcription regulation cascades projections represent an ideal framework for further developments.

## MATERIALS AND METHODS

### Software overview and implementation

SCITRAM (Single Cell Integrated Transcriptional Regulation Modeler), is a desktop graphical application developed in Python (3.10.11) for analysis and interpretation of single-cell RNA-sequencing data. The software provides data import, quality control, normalization, dimensionality reduction, cell clustering, differential-expression analysis, marker-based visualization and context-specific gene-regulatory-network reconstruction. The interface was implemented with Tkinter. Single-cell objects and preprocessing were handled with AnnData and Scanpy. Numerical, tabular and statistical operations within SCITRAM were implemented with NumPy, pandas, SciPy and statsmodels. Graph analysis is performed with NetworkX, and visualization with Matplotlib, Scanpy and Cytoscape.js. Windows executable was assembled with PyInstaller (6.17.0). Core package versions are listed below, and the complete environment is recorded in SCITRAM_requirements.txt. available at GitHub (https://github.com/SysFate/SCITRAM).

### Software environment

The following package versions were taken from the SCITRAM build environment; the complete transitive dependency list remains in the accompanying freeze file.

| **Package** | **Version** | **Role in SCITRAM** |
| --- | --- | --- |
| Python | 3.10.11 | Runtime used for the Windows build |
| Scanpy | 1.11.5 | Preprocessing, Scrublet, PCA, neighbors, UMAP, Leiden and integration wrappers |
| AnnData | 0.11.4 | Annotated single-cell data container |

| Package | Version | Role in SCITRAM |
| --- | --- | --- |
| NumPy | 2.2.6 | Array operations and network randomization |
| SciPy | 1.15.3 | Sparse matrices, Mann-Whitney tests and hierarchical clustering |
| pandas | 2.3.3 | Tables and metadata |
| scikit-learn | 1.7.2 | Machine-learning and neighbor dependencies |
| umap-learn | 0.5.9.post2 | UMAP implementation |
| igraph / leidenalg | 1.0.0 / 0.11.0 | Leiden graph clustering |
| harmonypy | 0.0.10 | Optional Harmony integration |
| bbknn | 1.6.0 | Optional batch-balanced neighbor graph |
| scanorama | 1.7.4 | Optional Scanorama integration |
| NetworkX | 3.4.2 | Regulatory-network analysis |
| statsmodels | 0.14.6 | FDR correction and network z-tests |
| Matplotlib / seaborn | 3.10.8 / 0.13.2 | Visualization |
| h5py | 3.15.1 | HDF5 input/output |
| pandastable | 0.14.0 | Interactive tables |
| PyInstaller | 6.17.0 | Windows executable packaging |

### Input data and data representation

SCITRAM accepts 10x Genomics HDF5 files, delimited count matrices (CSV, TSV or TXT), and 10x-style Matrix Market directories containing matrix.mtx, barcodes.tsv and features.tsv or genes.tsv, including gzip-compressed variants. Delimited matrices are imported as genes by cells and transposed to the AnnData cells-by-genes orientation. Matrix Market inputs are likewise transposed from genes by cells, and gene names are made unique. Each cell is annotated with its source sample in orig_file. Multiple objects are concatenated using an outer gene join and index_unique=’-’, which appends the sample key to cell identifiers.

Before normalization, the input count matrix is retained as a sparse CSR matrix in “adata.layers[’counts’]”. Non-finite values are replaced by zero, and negative values are rejected by integrity checks. This layer remains the authoritative source for cell- and gene-level count summaries, fold-change calculations, the fast differential-expression route, gene-expression overlays and count-based gene heatmaps. The working “adata.X” matrix is normalized to a total of 10,000 counts per cell and optionally log-transformed. A copy of this normalized, unscaled expression matrix is stored in “adata.layers[’norm’]” before optional scaling or cell-cycle regression. The resulting working matrix is used for highly variable gene selection, principal-component analysis, optional batch integration and neighborhood-graph construction, and is cleared after clustering to reduce memory use. Cluster-specific lightweight “AnnData” objects retain raw counts in “layers[’counts’]” and normalized expression in “X”, together with selected observation metadata, UMAP coordinates, and per-gene raw-count totals and means. Normalized cluster expression is used for mean-expression summaries and marker-category heatmaps, whereas raw counts remain the source for count-derived percentages and fold-change calculations.

### Quality control and filtering

For multiple HDF5, Matrix Market, and delimited inputs, filtering is applied independently to each sample before concatenation. An optional random subsample can first be drawn as a user-defined percentage of cells. Genes are retained according to a minimum number of expressing cells, and cells are sequentially filtered by total UMI counts, detected genes and mitochondrial transcript percentage using Scanpy. Mitochondrial genes are detected case-insensitively from the “MT-” prefix, and cells are retained when “pct_counts_mt” is strictly below the selected threshold.

The graphical-interface defaults are genes detected in at least 3 cells; 500-15,000 UMI counts per cell; 200-25,000 detected genes per cell; <5% mitochondrial counts; and 100% of cells retained during optional subsampling. Thresholds are user adjustable. The QC report records settings, sample-level before/after counts for each filter, blacklist status, mitochondrial summaries, doublet results, normalization settings and dimensionality-reduction parameters.

An optional blacklist removes genes listed in bundled Ensembl release 115 human or mouse tables. Human and mouse selection is inferred from the presence of “MT-” and “Mt-” gene prefixes, respectively, with human used as the fallback. The distributed tables (Human or mouse “_blacklist_genes_Ensembl115.tsv” in “ExtraFiles” directory) identify genes through an “hgnc_symbol” column and include mitochondrial, immunoglobulin, major-histocompatibility-complex, ribosomal and hemoglobin-related categories.

### Doublet detection

When enabled (default), doublets are detected with Scanpy’s Scrublet wrapper using “Princial components = 20” and “Neighbors = 10”. For datasets containing more than one origin file category, detection is stratified with “batch_key=’orig_file’”. Cells labeled “predicted_doublet” are removed. If Scrublet fails, the exception and failed status are recorded, and analysis continues without doublet removal.

### Normalization and optional covariate regression

Library-size normalization is performed using “scanpy.pp.normalize_total” with “target_sum=10,000” counts per cell, followed by an optional “log1p” transformation that is enabled by default. Following normalization and log transformation, a copy of the normalized, unscaled expression matrix is stored in “adata.layers[’norm’]” for downstream mean-expression summaries and marker-category heatmaps. Optional scaling is subsequently performed using “scanpy.pp.scale”, with zero-centering controlled independently by the user; both scaling and zero-centering are disabled by default. Optional cell-cycle regression is applied only to the working “adata.X” matrix. Non-finite processed values are replaced by zero, whereas the raw-count matrix stored in “adata.layers[’counts’]” remains unchanged. The resulting working “adata.X” matrix is used for highly variable gene selection and principal-component analysis, while “adata.layers[’norm’]” retains normalized expression values that are not affected by subsequent scaling or regression.

Cell-cycle effects can optionally be regressed out. Built-in human or mouse S-phase and G2/M gene sets are used to calculate these scores with “scanpy.tl.score_genes_cell_cycle”; both scores are then regressed from the processed expression matrix with “scanpy.pp.regress_out” using five processing cores by default. Inferring species for this step came from the first gene name, lowercase for mouse and capitals for human.

### Highly variable genes, dimensionality reduction and clustering

Highly variable genes are identified with “scanpy.pp.highly_variable_genes” using a user-selected “top genes” value (2,000 by default); the call also supplies min_mean=0.05, max_mean=1.5 and min_disp=0.5. Principal-component analysis is performed with “scanpy.tl.pca” using the “ARPACK solver” and 20 components by default. Batch correction is optional and defaults to None. When requested for a dataset with at least two origin files categories, Harmony stores an adjusted PCA representation in “X_pca_harmony”, BBKNN directly constructs a batch-balanced neighbor graph, and Scanorama stores an integrated representation in “X_scanorama”. Harmony and Scanorama representations are used for subsequent neighbor construction; BBKNN replaces the ordinary neighbor step. Raw counts remain unchanged. Requests made for a single sample are skipped, whereas missing optional dependencies or integration errors stop dimensional analysis rather than silently falling back to an uncorrected result.

Without batch correction (by default), a neighborhood graph is calculated with “scanpy.pp.neighbors” using 15 nearest neighbors and 20 principal components by default. For BBKNN, neighbors_within_batch is set to “max(1, ceil(n_neighbors/n_batches))”. UMAP coordinates are generated with “scanpy.tl.umap”. Cell communities are identified with “scanpy.tl.leiden” at a default resolution of 0.5 and random_state=0. Technical cluster identifiers are relabeled consecutively from 1 for presentation.

SCITRAM also supports local reclustering of a selected parent cluster. Raw counts are normalized to a total of 10,000 counts per cell and log-transformed, genes detected in fewer than one cell are removed, and highly variable genes are selected using Scanpy’s “Cell Ranger dispersion-based” method. PCA, neighborhood-graph construction, UMAP and Leiden clustering are subsequently performed using user-defined parameters. Resulting subclusters receive “parent_child” identifiers.

### Cluster-level expression summaries and fold changes

For every Leiden cluster, SCITRAM calculates the raw-count mean and total for each gene across all cells assigned to that cluster. For the cluster-versus-rest summary, the fold change for gene g in cluster c is calculated as:

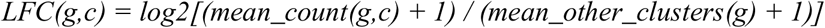

where mean_other_clusters is the unweighted mean of the cluster-level mean counts across all clusters other than c, and 1 is the pseudocount. For comparisons involving one or more target and reference clusters, raw-count matrices are pooled at the cell level for statistical testing; for network analysis, the cluster-level gene means are first averaged within the selected target and reference cluster sets and the same pseudocount-based log2 ratio is calculated.

### Differential-expression analysis

The fast target-versus-reference comparison operates directly on the preserved raw-count matrices. Cells from all selected target clusters and all selected reference clusters are stacked within their respective groups. For each gene, SCITRAM reports the arithmetic mean raw count and percentage of cells with a non-zero count in each group. Log2 fold change is calculated as:

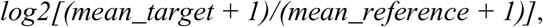

and average abundance for MA-type displays is calculated as:

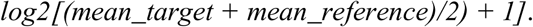

A two-sided Mann–Whitney U test is applied independently to each gene using the asymptotic method. Genes with zero counts in both groups are assigned P = 1. P values are adjusted across all tested genes using the Benjamini–Hochberg false-discovery-rate procedure. The reported score is the absolute standardized U statistic:

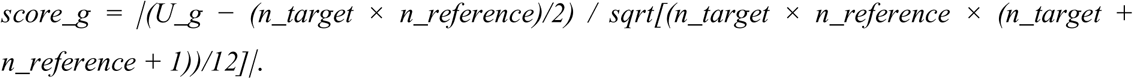

Results can be filtered by user-defined absolute log2-fold-change and adjusted-P-value thresholds and ranked for tabular or graphical output. For cluster-marker analysis, SCITRAM additionally exposes “scanpy.tl.rank_genes_groups*”* with the Wilcoxon method and Benjamini-Hochberg correction.

### Gene-expression visualization and heatmaps

Expression of individual genes or user-selected gene sets can be projected onto the existing UMAP using raw counts. For selected gene lists, counts are aggregated per cell before color mapping. Cluster summaries report both total counts for the selected genes and counts divided by the number of cells in each cluster.

For gene-level heatmaps, the selected raw-count columns are extracted across all cells and standardized gene-wise with “scanpy.pp.scale” using zero centering and “max_value = 2”. Cluster-level average standardized expression is displayed with “scanpy.pl.matrixplot”, using Leiden clusters as groups, hierarchical clustering of clusters, a blue–white–red color scale and a displayed range of −2 to 2.

For marker-category heatmaps, genes are grouped by a user-supplied “marker type” annotation list. Within each cell cluster, values are averaged across all cells and available genes assigned to a marker category. Each marker-category row is converted to a Z score across clusters, with the standard deviation bounded below by 10^-6, and values are clipped to [-2, 2]. Rows and columns are hierarchically clustered using Euclidean distance and average linkage.

### Regulatory-network reconstruction

SCITRAM reconstructs a condition-specific transcriptional regulatory network by integrating a prior gene-regulatory-network (GRN) collection (CellNet_[ Human or Mouse]_GRN032014_PTOX.csv files distributed in “GRN-collection” directory ^7^) with expression changes calculated between user-selected target and reference cell clusters. The prior network is represented as a directed edge table containing a transcription factor (TF), target gene (TG), and one or more edge-confidence or correlation columns. The user should select the GRN, the edge-associated correlation column, a minimum absolute correlation threshold, target and reference clusters, and positive and negative log2-fold-change thresholds.

### GRN filtering and coherent edge selection

Target- and reference-group gene means are derived from raw counts. When several clusters are selected in either group, cluster means are averaged with equal weight. Gene-wise log2 fold changes are computed with a pseudocount of 1 and joined to the prior GRN by target-gene identity; targets missing from the expression matrix are assigned LFC = 0. Prior edges are first retained when the absolute value of the selected correlation metric is at least the user-defined minimum. A coherent directed edge is then defined by agreement between the sign of the TF–TG association and the direction of target-gene change: edges with corr ≥ 0 require LFC ≥ 0, whereas edges with corr < 0 require LFC < 0.

By default, propagation is restricted to edges whose target genes exceed the selected positive or negative LFC thresholds. An optional setting retains the complete coherent network, including non-responsive target genes. The default network thresholds are LFC > 0.5 or LFC < −0.5 and |corr| ≥ 0; all are user-adjustable.

### Propagation yield and topological statistics

The coherent network is converted to a directed “NetworkX” graph. In the current implementation, all TF nodes present as edge sources in the propagation network are evaluated as starting nodes. For each starting TF, the reachable subgraph consists of the TF itself and all downstream descendants. Propagation yield is defined as 100 × (number of nodes in the reachable subgraph)/(total number of nodes in the propagation graph). Equivalent yields are also calculated after restricting breadth-first traversal to depths 1, 2 and 3 (optional). Network topology was represented using a directed NetworkX graph (DiGraph), with regulatory edges oriented from transcription factors to their target genes (TF→TG). Directed betweenness centrality was calculated using NetworkX 3.4.2 with betweenness_centrality(normalized=True). The calculation considered all directed shortest paths, excluded path endpoints and treated every retained edge as having unit length; consequently, correlation and fold-change magnitudes were not used as path weights. For each node (v), normalized betweenness centrality was defined as

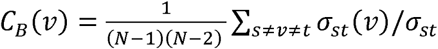

where (*σ_st_*) is the number of directed shortest paths between nodes (s) and (t), and (*σ_st_*(*v*)) is the number of those paths passing through (v). Total node degree was calculated as the sum of the unweighted in-degree and out-degree. SCITRAM additionally defined a bottleneck score as the product of normalized betweenness centrality and total node degree.

### Network randomization and statistical confidence

To establish a null distribution for propagation yield, SCITRAM independently permutes the correlation values among the retained propagation-network edges. After each permutation, the coherence rule between the randomized correlation sign and the observed target-gene LFC is reapplied, a randomized directed graph is constructed, and propagation yield is recalculated for every starting TF. The default interface performs 20 iterations. For each TF, SCITRAM records the mean, standard deviation and 10th, 90th and 95th percentiles of randomized yields.

The implementation currently reports a one-sided parametric z-test by calling “statsmodels.stats.weightstats.ztest” on the randomized yields with the observed yield as the null value and alternative = ‘smaller’. If the randomized standard deviation is zero, P is set to 1. Network P values are presently reported without multiple-testing correction. Delta yield is calculated as observed yield − (mean randomized yield + one standard deviation).

### Influence score and ranking

A distance-weighted influence score combines propagation at the first three network depths. With α = 0.7, the base score is exp(−α)·yield_d1 + exp(−2α)·yield_d2 + exp(−3α)·yield_d3. In the active thresholded weighting mode, this value is multiplied by max(|LFC| − LFC_threshold, 0), where LFC_threshold is the larger absolute value of the positive and negative thresholds. Scores are linearly rescaled so that the maximum equals 100. TFs are ranked sequentially by influence score, delta yield and total propagation yield. Interactive subnetworks and edge/node tables can be exported for downstream exploration.

### Automatic LFC-threshold exploration

SCITRAM can evaluate a user-defined series of symmetric LFC thresholds (default range 0.1–1.0 in increments of 0.1). For each threshold, the software rebuilds the coherent, LFC-restricted propagation network and reports the number of responsive genes, nodes, edges and weakly connected components, together with directed graph density E/[N(N−1)]. This exploration is topological only and does not calculate propagation yields or randomized significance until a threshold is selected for full analysis.

### Reproducibility and outputs

SCITRAM records analysis date, input samples, filtering histories, doublet-removal results, normalization settings and dimensionality-reduction parameters in a quality-control report stored under scitram_qc during the live analysis and copied into the saved project metadata. Cell-level QC metrics are exported as timestamped CSV files. A saved project is packaged as a ZIP containing a lightweight H5AD file and a pickle metadata file. The H5AD retains observation and variable metadata, layers[’counts’] and X_umap; the pickle retains cluster objects, colors, QC information and plot settings. PCA coordinates, neighbor graphs and batch-corrected representations are not currently retained in the lightweight H5AD, so a saved project preserves the obtained clustering and visualization but is not, by itself, a complete object for reconstructing global clustering. Graphical outputs, including global and local UMAP representations, differential-expression plots, gene- and marker-category heatmaps, and ranked-transcription-factor plots, can be exported directly from the graphical interface as 300-dpi PNG images or as vector PDF and SVG files. Gene-expression violin plots are currently exported as 300-dpi PNG images.

Gene regulatory networks displayed in figures 3, 4 and 5 were produced in Cytoscape by using the exported network format produced by SCITRAM.

## SUPPLEMENTAL INFORMATION

Supplementary Figures S1-S9; Supplementary Table S1.

## ACKNOWLEDGMENTS

We thank all members of the European consortium T-FITNESS as well as the team members of SysFate for contributing to the discussion of this project.

## AUTHOR CONTRIBUTIONS

Ariel Galindo (Investigation, Data curation, Software development, Formal analysis), Maximilien Duvina (Formal analysis, Software development, Data curation), Paula Barbao (Resources, Methodology), Alba Rodriguez-Garcia (Resources, Methodology), Christoph Heuser (Resources, revision original draft), Sonia Guedan (Supervision, Funding acquisition, revision original draft), Luca Gattinoni (Supervision, Funding acquisition, revision original draft) and Marco Antonio Mendoza-Parra (Conceptualization, Formal analysis, Supervision, Funding acquisition, Writing original draft).

## DATA AND SOFTWARE AVAILABILITY

SCITRAM is available at GitHub (https://github.com/SysFate/SCITRAM).

CD8+ Single cell transcriptomics data related to the PanCancer study were obtained from GSE15678 (ThyroidCa_THCA, Endometrial_UCEC, RenalCa_RC, EsophangealCa_ESCA, MultipleMyolome_MM, PancreaticCa_PACA, BC, BCellLy_BCL, GSM4743199_OvarianCa_OV, GSM4743231_FallopianCa_FTC, GSM4743237_CholangioCa_CHOL_SS2).

CD8+ single cell transcriptomics data related to the infiltrating T cells isolated from SKOV3 tumors were obtained within the context of the T-FITNESS consortium (single-cell RNA sequencing are available in GEO: GSE319746).

Pancreatic endocrinogeneis data were obtained from RegVelo repository: https://figshare.com/articles/dataset/Pancreatic_endocrinogenesis/28124579. Downloaded .h5ad files were converted in MTX format compatible with SCITRAM with an in-house python script.

Human brain organoids data has been obtained from the repository https://www.synapse.org (accession number project ID syn26346373). Seurat objects obtained for each time-point were converted in matrix file format compatible with SCITRAM by using an in-house R script.

All files compatible with SCITRAM used in this study are available in https://figshare.com/account/articles/33977794

