## Supplementary_Figures for "SCITRAM: a Single-cell Integrated transcription regulation modeler for predicting master transcription factors"

#### Supplementary Material

**Supplementary Figure S1.** Heatmap displaying the normalized z-score gene expression per cluster associated to the effector or the exhausted state retrieved within the pan-cancer single-cell atlas.

**Supplementary Figure S2.** Gene co-regulatory networks assessed from master TFs predicted by SCITRAM at different Log2 fold-change (LFC) thresholds in the universal pancancer cell Atlas.

**Supplementary Figure S3.** Number of common and specific differentially expressed genes between T cell clusters issued from day 27 (clusters 3, 4 and 7) relative to day 8 (cluster 2) after CAR-T cell treatment of solid tumors.

**Supplementary Figure S4.** Gene regulatory network (GRN) characteristics associated to the indicated cluster comparisons.

**Supplementary Figure S5.** Heatmap displaying the normalized z-score gene expression per cluster associated to the different cell types composing the pancreatic cell lineage.

**Supplementary Figure S6.** Gene regulatory network (GRN) characteristics associated to the the analysis of the indicated pancreatic cell types.

**Supplementary Figure S7.** Master TF co-regulatory network assessed for the indicated pancreatic cell types.

**Supplementary Figure S8.** Mean gene expression enrichment levels associated to the inferred pancreatic cell fate regulomes.

**Supplementary Figure S9.** Venn Diagrams displaying the common and specific master transcription factors revealed by SCITRAM for each of the indicated clusters during cerebral organoids differentiation.

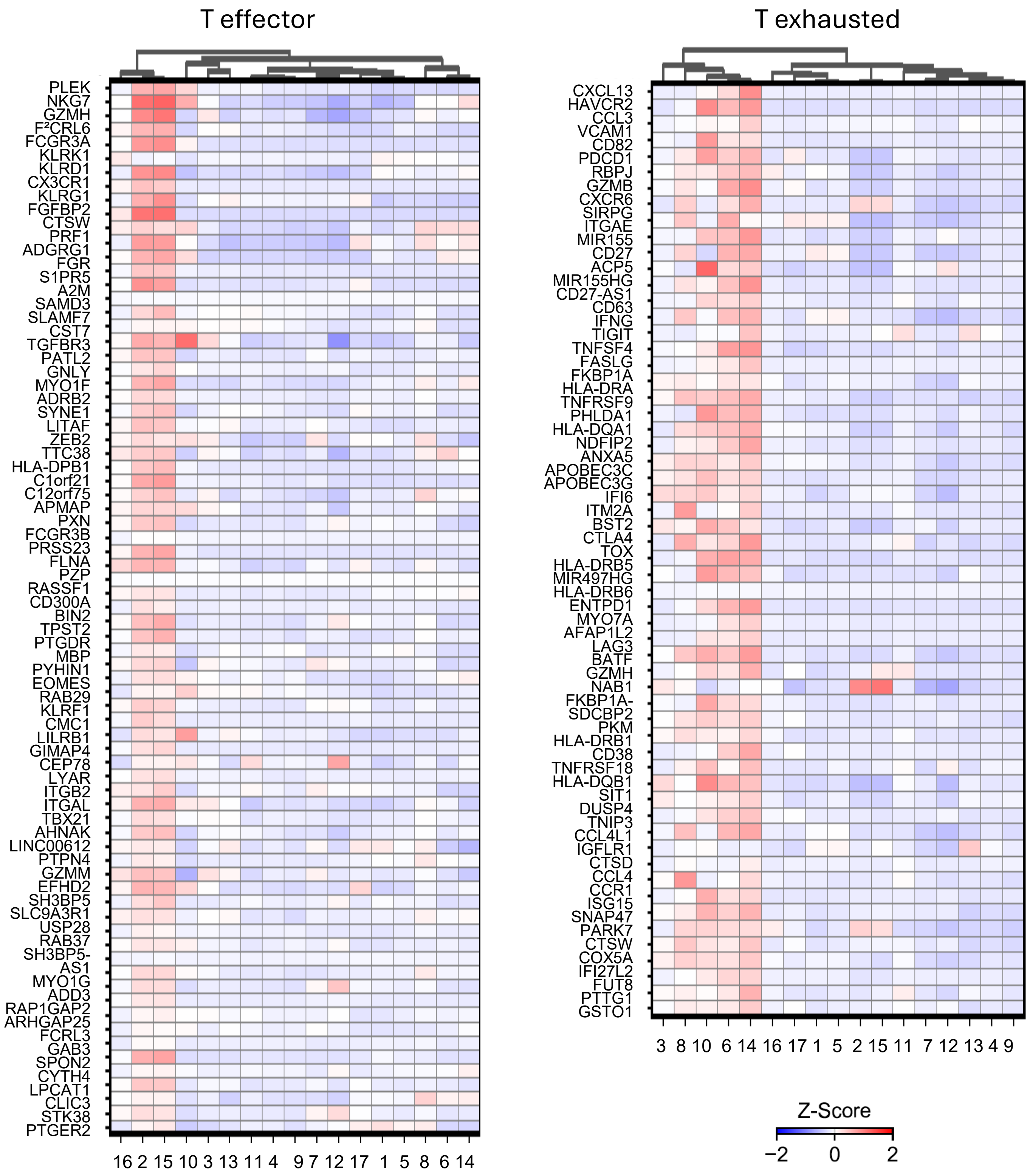


**Supplementary Figure S1. Heatmap displaying the normalized z-score gene expression per cluster associated to the effector or the exhausted state retrieved within the pan-cancer single-cell atlas.**

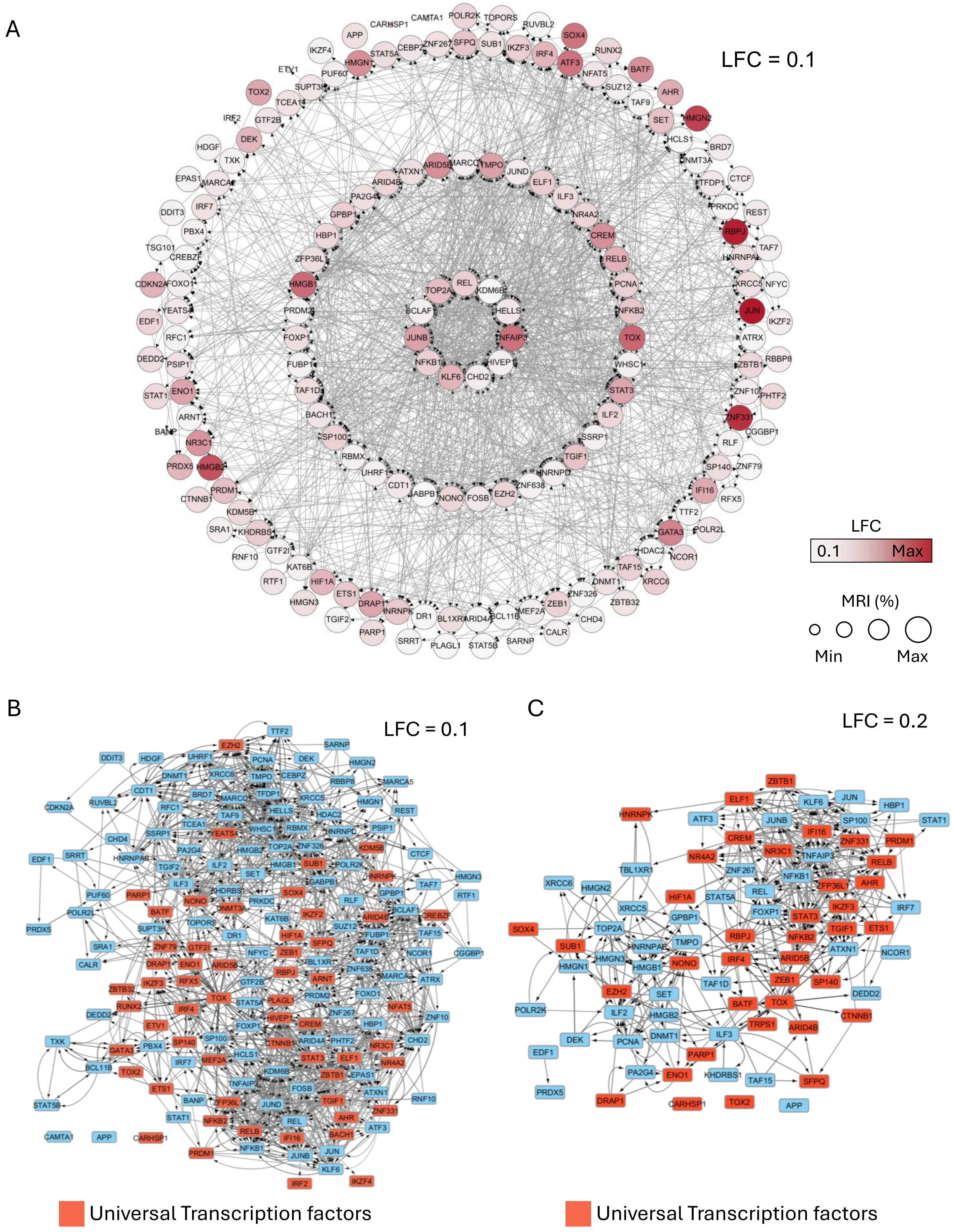


**Supplementary Figure S2. Gene co-regulatory networks assessed from master TFs predicted by SCITRAM at different Log2 fold-change (LFC) thresholds in the universal pancancer cell Atlas. (A)** GRN assessed for a LFC threshold of 0.1. (B) Same as (A) where TFs defined as part of the universal exhaustion list by Zheng et al (Science 2021) are depicted in red. (B) TF co-regulatory network assessed for a LFC threshold of 0.2.

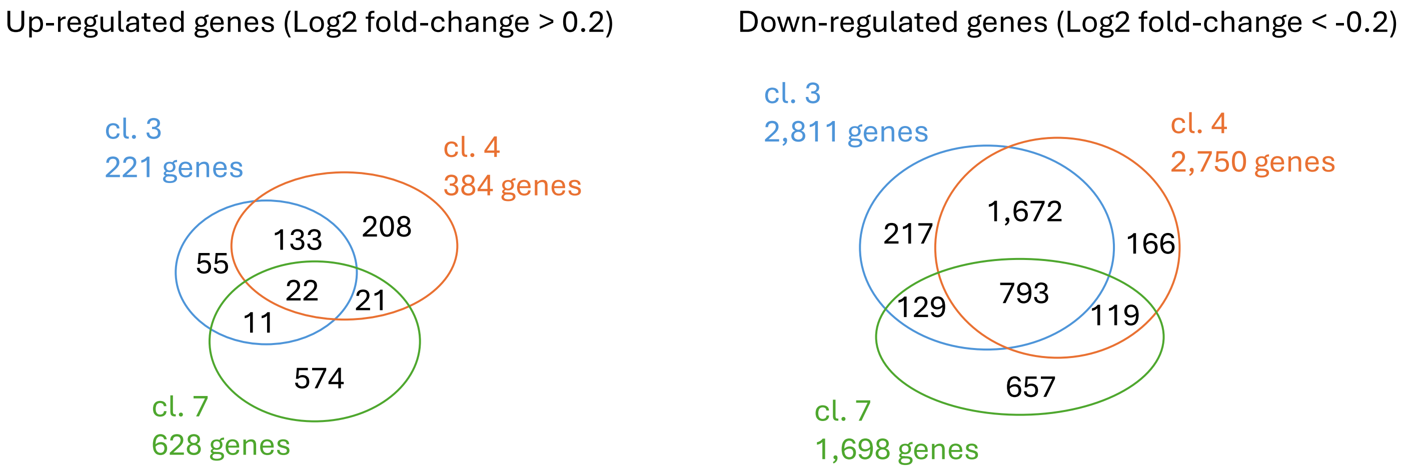


**Supplementary Figure S3. Number of common and specific differentially expressed genes between T cell clusters issued from day 27 (clusters 3, 4 and 7) relative to day 8 (cluster 2) after CAR-T cell treatment of solid tumors.**

**Cl. 3 vs cl. 2**

| LFC | DEG | Nodes | Edges |
|---|---|---|---|
| 0.1 | 5560 | 5971 | 45893 |
| 0.2 | 3053 | 3884 | 26150 |
| 0.3 | 1720 | 2754 | 15207 |
| 0.4 | 1008 | 2086 | 9257 |
| 0.5 | 585 | 1568 | 5634 |
| 0.6 | 363 | 1231 | 3657 |
| 0.7 | 216 | 938 | 2257 |
| 0.8 | 123 | 664 | 1325 |
| 0.9 | 80 | 544 | 950 |
| 1 | 52 | 421 | 654 |

**Cl. 4 vs cl. 2**

| LFC | DEG | Nodes | Edges |
|---|---|---|---|
| 0.1 | 5469 | 5880 | 47978 |
| 0.2 | 3044 | 3925 | 28135 |
| 0.3 | 1757 | 2813 | 16995 |
| 0.4 | 1064 | 2177 | 11048 |
| 0.5 | 663 | 1734 | 7009 |
| 0.6 | 411 | 1374 | 4395 |
| 0.7 | 263 | 1101 | 2921 |
| 0.8 | 166 | 852 | 1827 |
| 0.9 | 108 | 692 | 1343 |
| 1 | 72 | 552 | 951 |

**Cl. 7 vs cl. 2**

| LFC | DEG | Nodes | Edges |
|---|---|---|---|
| 0.1 | 5993 | 6283 | 50951 |
| 0.2 | 3585 | 4359 | 30593 |
| 0.3 | 2305 | 3274 | 19375 |
| 0.4 | 1442 | 2490 | 12143 |
| 0.5 | 893 | 1937 | 7624 |
| 0.6 | 577 | 1548 | 4916 |
| 0.7 | 382 | 1225 | 3203 |
| 0.8 | 241 | 944 | 2007 |
| 0.9 | 158 | 729 | 1345 |
| 1 | 113 | 581 | 986 |

**Supplementary Figure S4. Gene regulatory network (GRN) characteristics associated to the indicated cluster comparisons.** LFC: Log2 Fold-change threshold used for defining the genes thar participate at the GRN reconstruction. DEG: Differentially expressed genes obtained with the indicated LFC. Nodes: Number of genes composing the GRN. Edges: Number of gene inter-relationships (connections) retrieved in the GRN.

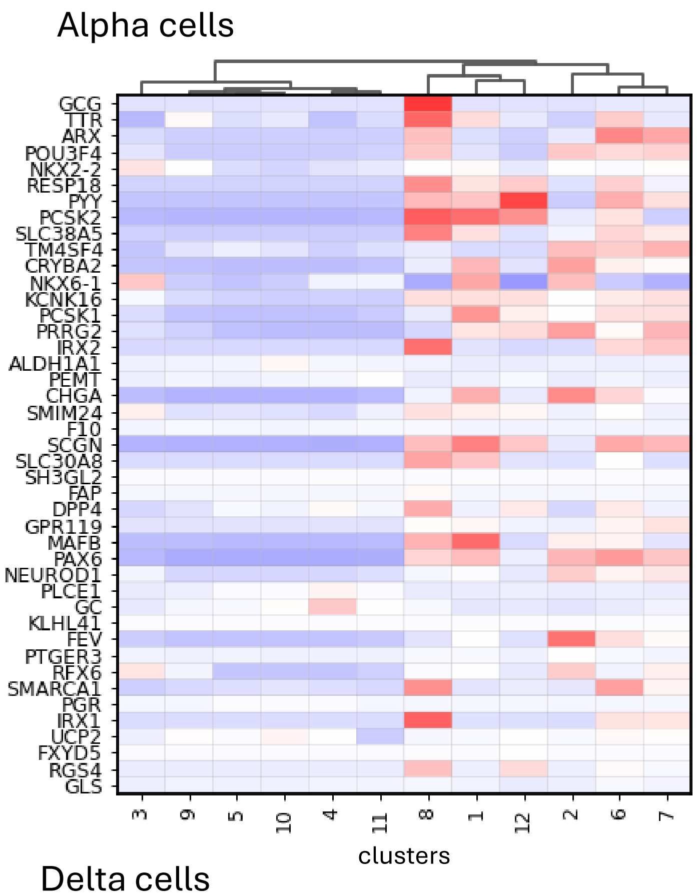


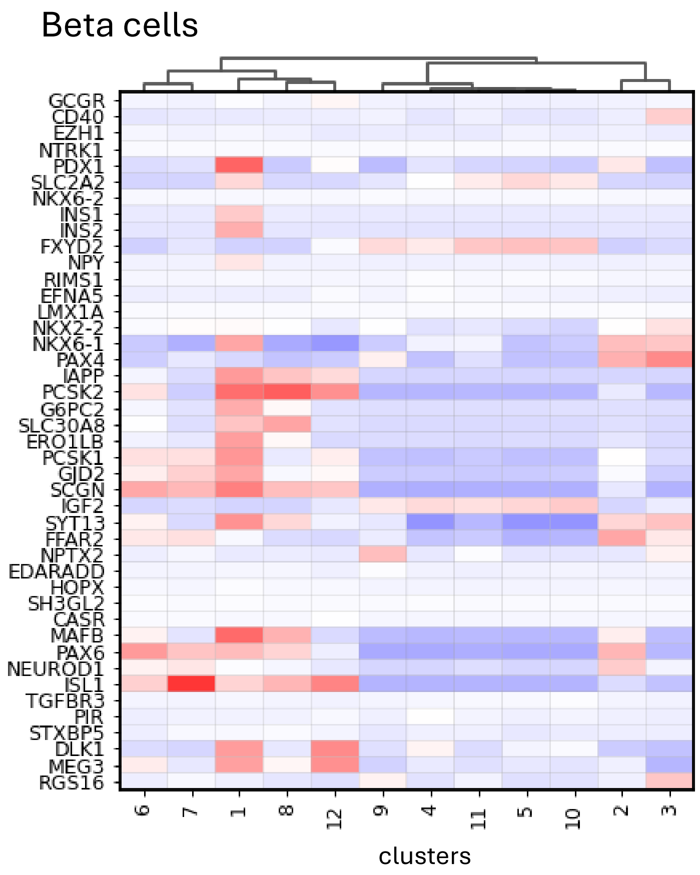


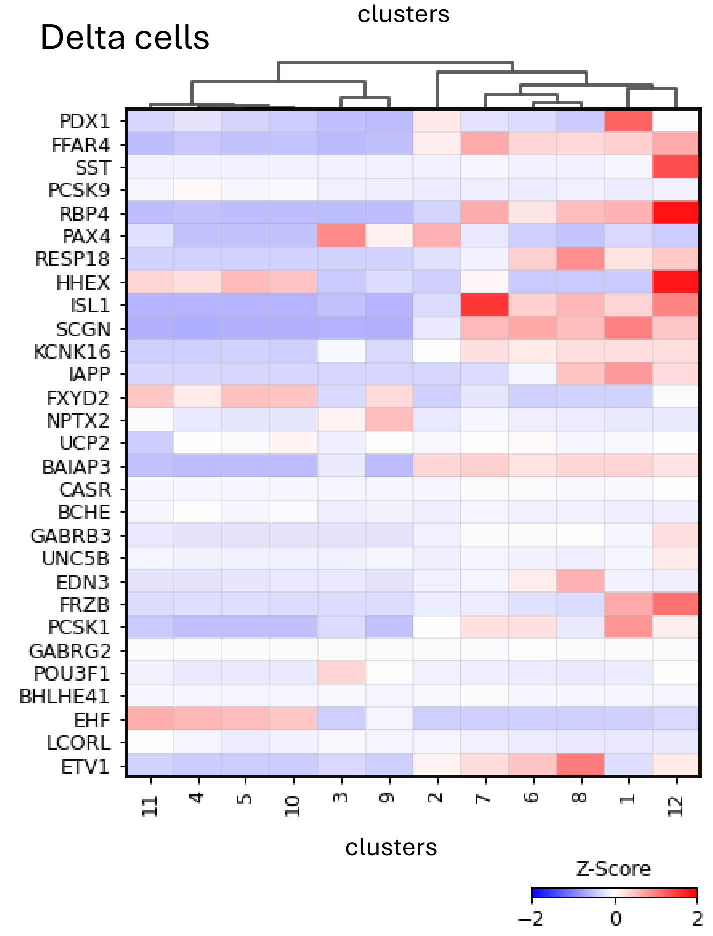


**Supplementary Figure S5. Heatmap displaying the normalized z-score gene expression per cluster associated to the different cell types composing the pancreatic cell lineage.**

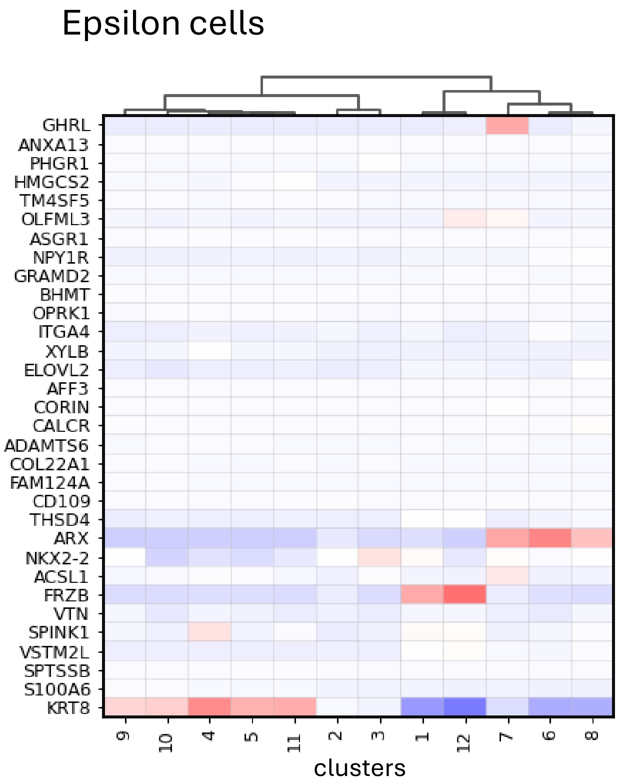


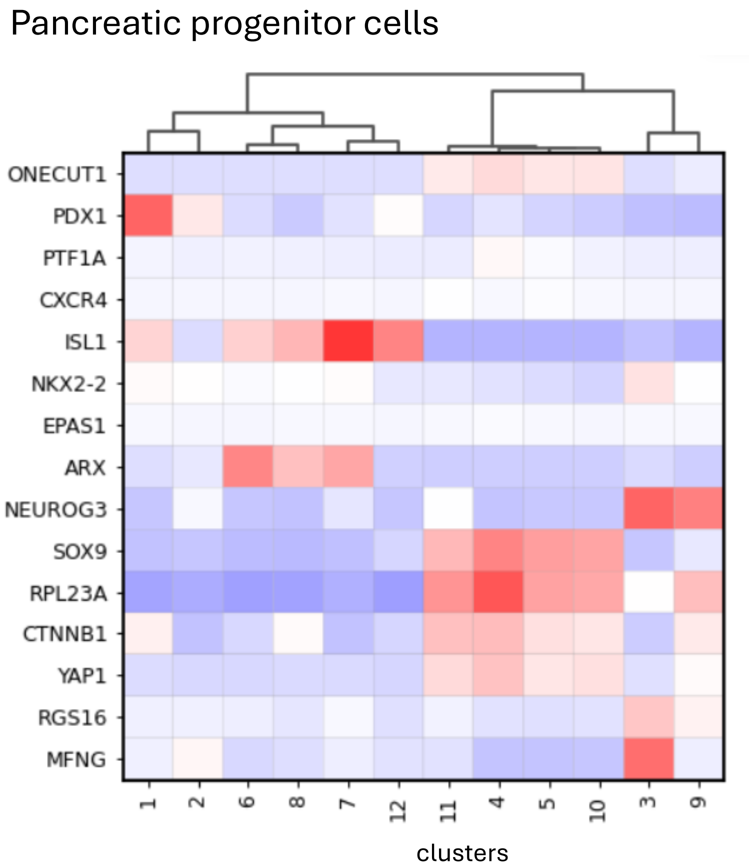


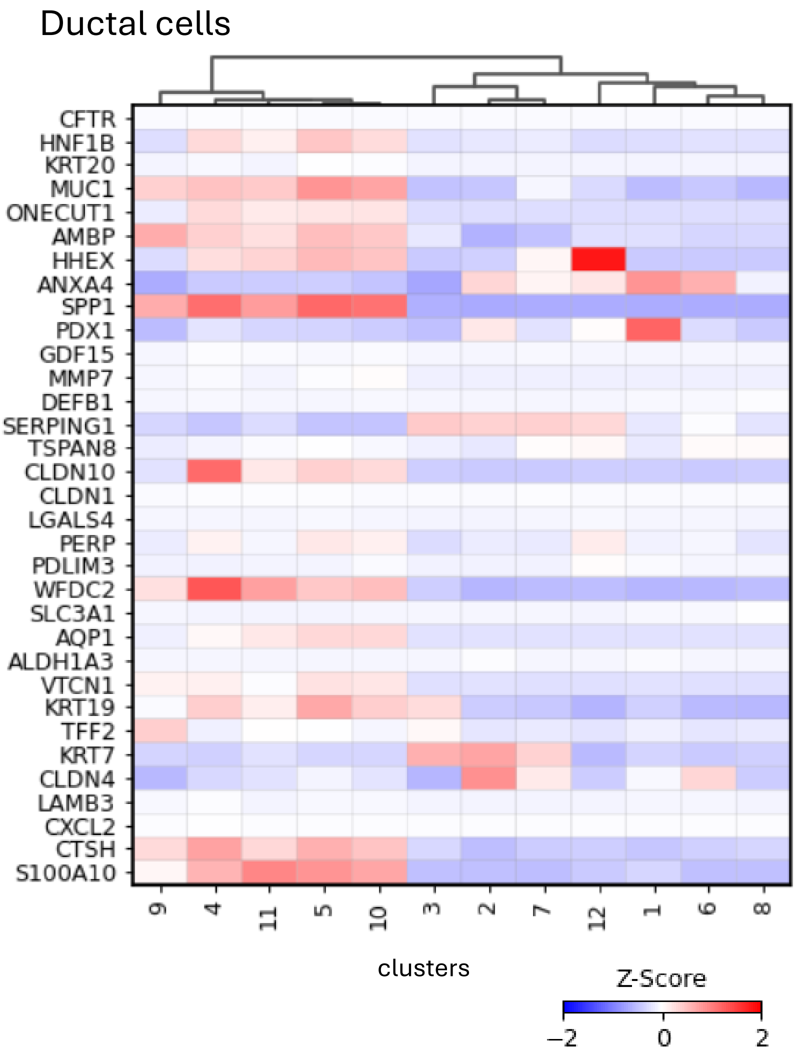


**Supplementary Figure S5. Heatmap displaying the normalized z-score gene expression per cluster associated to the different cell types composing the pancreatic cell lineage.**

A

Log2 Fold-change filter: 0.1

| Cell type | clusters | DEG | Nodes | Edges |
|---|---|---|---|---|
| Progenitors | cl.3.cl.9 | 3,295 | 4,220 | 37,225 |
| Ductal | cl5.cl.10 | 3,309 | 4,229 | 41,930 |
| Alpha | cl.8 | 3,494 | 4,379 | 40,013 |
| Beta | cl.1 | 3,368 | 4,293 | 38,440 |
| Delta | cl.12 | 3,982 | 4,770 | 39,876 |
| Epsilon | cl.7 | 2,949 | 3,932 | 32,806 |

**Supplementary Figure S6. Gene regulatory network (GRN) characteristics associated to the the analysis of the indicated pancreatic cell types.** Clusters: cell clusters used for a differential expression analysis relative to all clusters. DEG: Differentially expressed genes obtained with the indicated LFC. Nodes: Number of genes composing the GRN. Edges: Number of gene inter-relationships (connections) retrieved in the GRN. The analysis has been performed for a log2 fold-change threshold of 0.1.

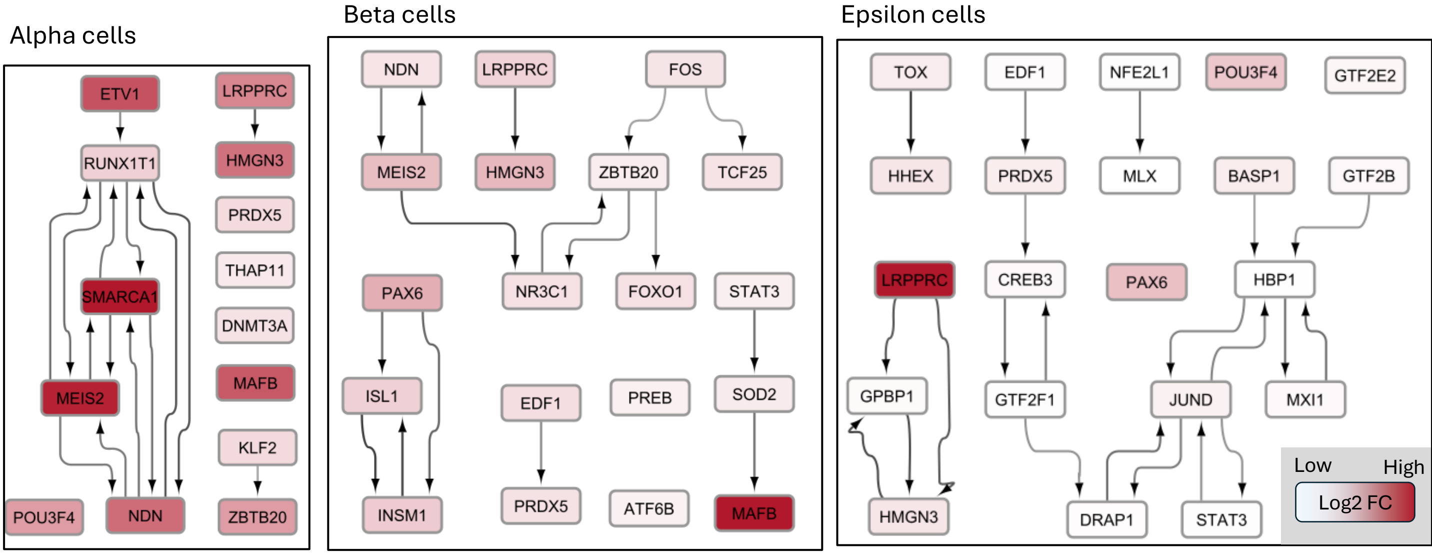


**Supplementary Figure S7. Master TF co-regulatory network assessed for the indicated pancreatic cell types.** Displayed GRNs were been performed for a log2 fold-change threshold of 0.2, as a way to reduce network complexity.

### Epsilon cells Regulome

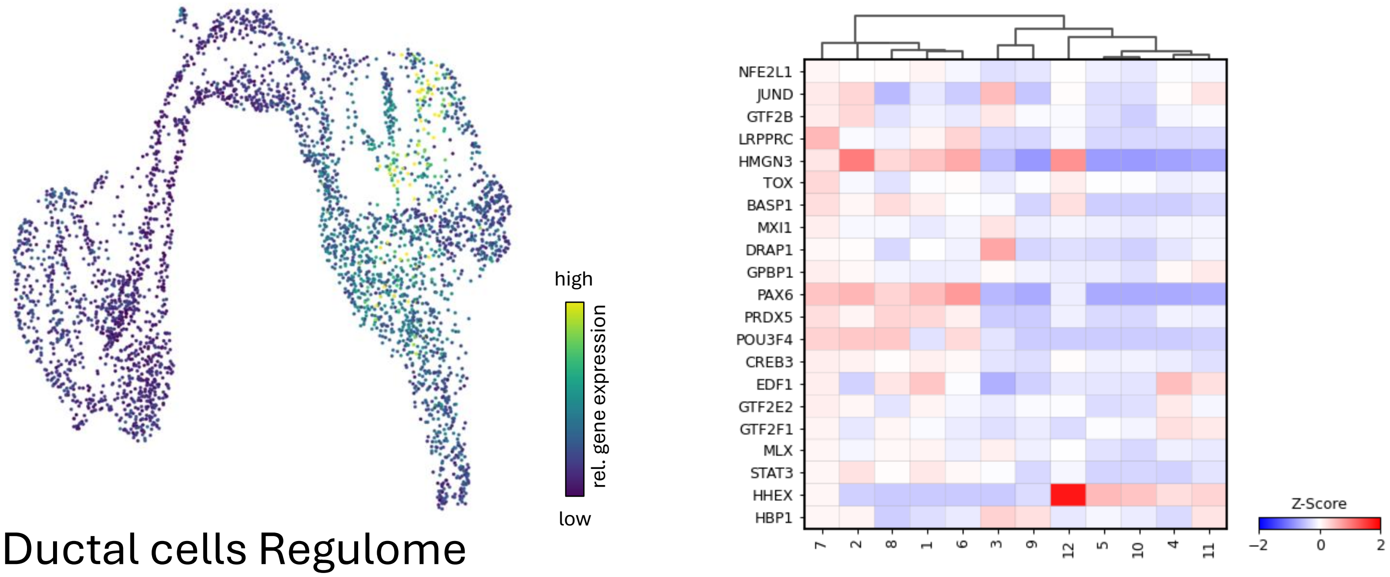


### Ductal cells Regulome

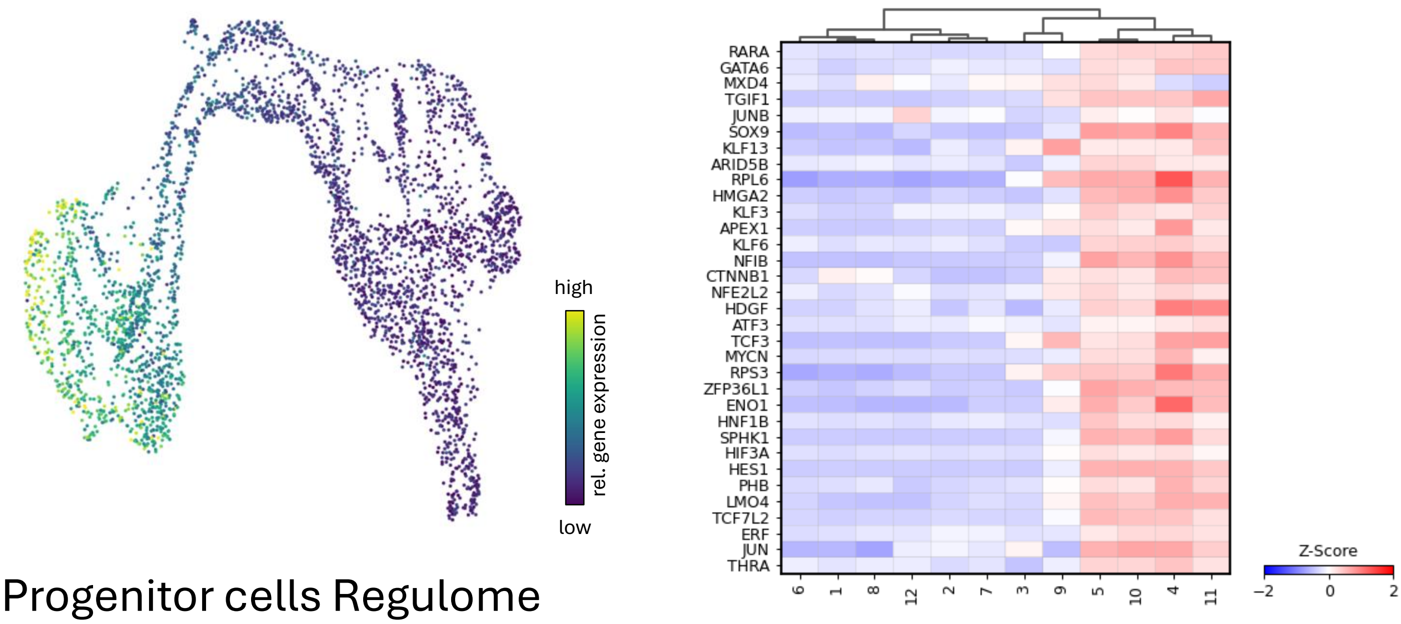


### Progenitor cells Regulome

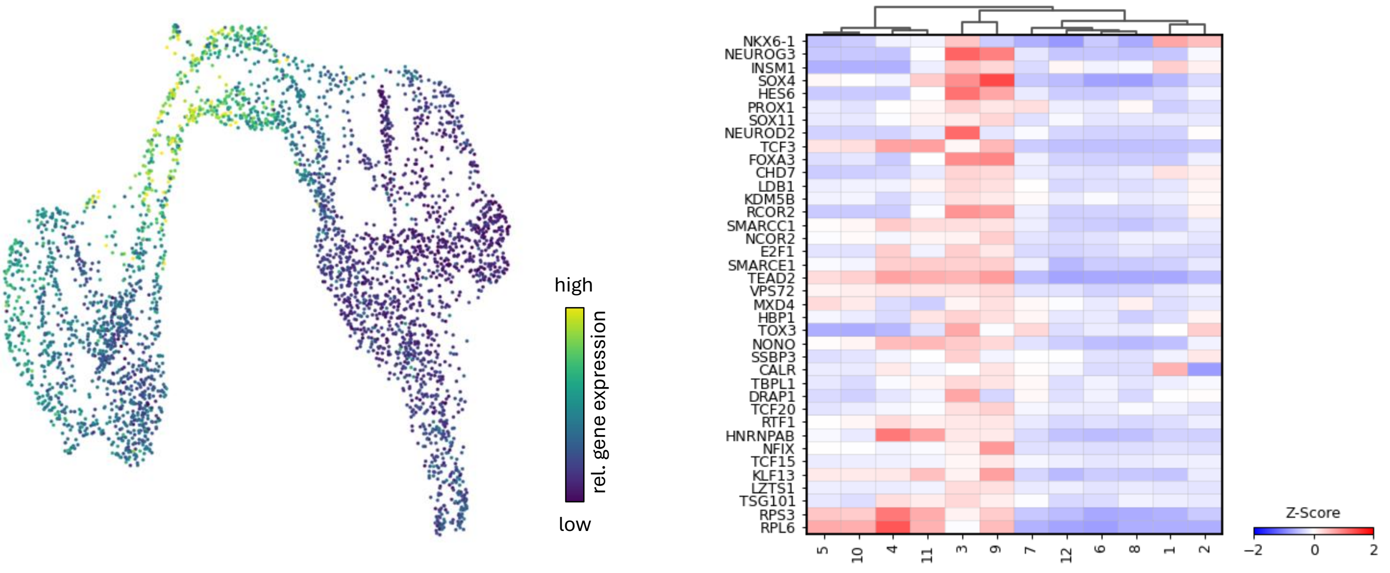


**Supplementary Figure S8. Mean gene expression enrichment levels associated to the inferred pancreatic cell fate regulomes.** Displays are produced by uploading to SCITRAM the top-ranked TFs associated to each pancreatic cell type. Left: Mean TF expression levels. Right: TF enrichment per clusters composing the UMAP.

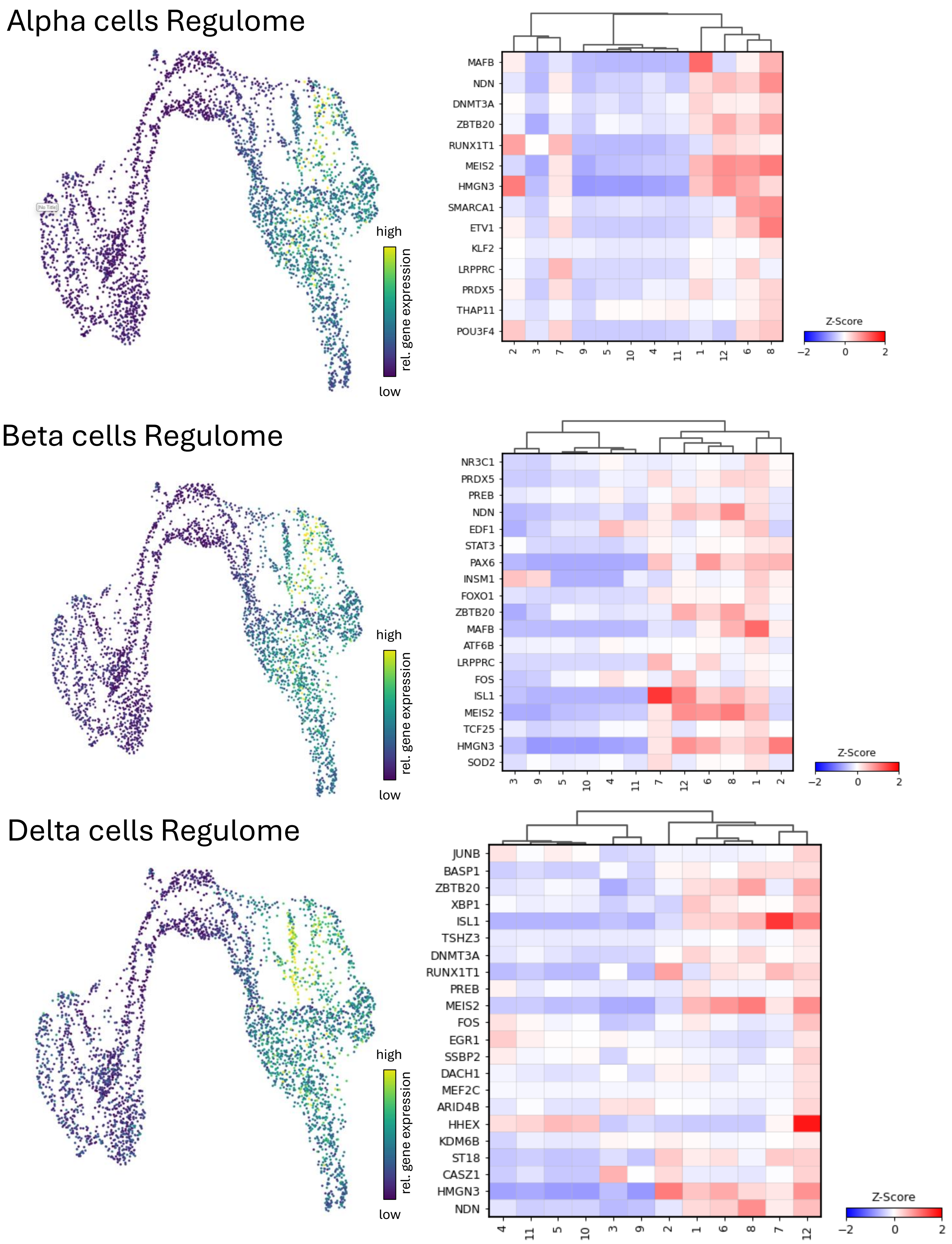


**Supplementary Figure S8. Mean gene expression enrichment levels associated to the inferred pancreatic cell fate regulomes.** Displays are produced by uploading to SCITRAM the top-ranked TFs associated to each pancreatic cell type. Left: Mean TF expression levels. Right: TF enrichment per clusters composing the UMAP.

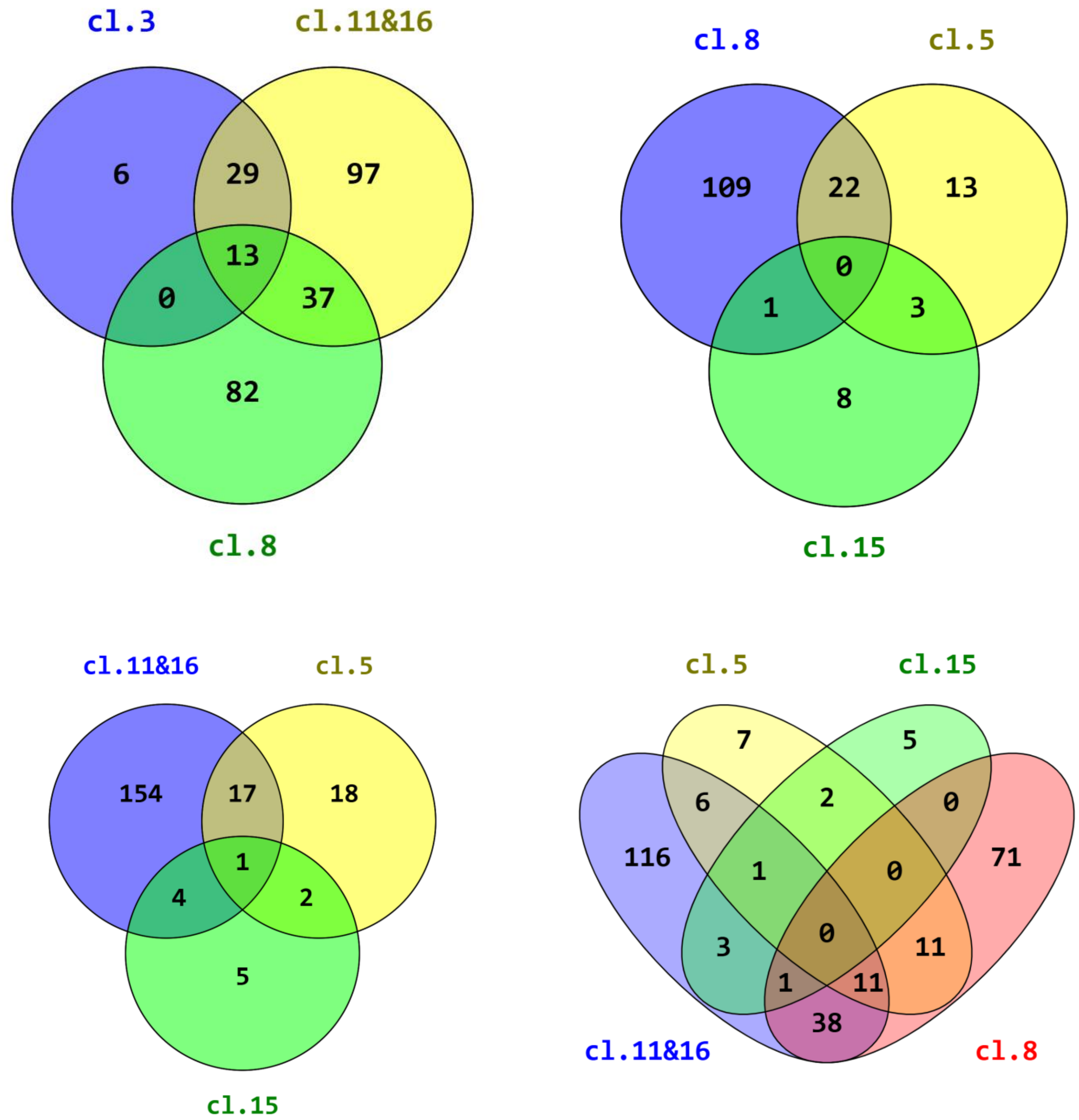


**Supplementary Figure S9. Venn Diagrams displaying the common and specific master transcription factors revealed by SCITRAM for each of the indicated clusters during cerebral organoids differentiation.**
